# Induction by caterpillars of stored and emitted volatiles in terpene chemotypes from populations of wild cotton (*Gossypium hirsutum*)

**DOI:** 10.1101/2024.04.17.589906

**Authors:** Marine Mamin, Mary V Clancy, Galien Flückiger, Teresa Quijano-Medina, Biiniza Pérez-Niño, Luis Abdala-Roberts, Ted C.J. Turlings, Carlos Bustos-Segura

## Abstract

**Background:** Upland cotton (*Gossypium hirsutum*) plants constitutively store volatile terpenes in their leaves, which are steadily emitted at low levels. Herbivory leads to a greater release of these stored volatiles. Additionally, damaged plants increase the accumulation of volatile terpenes in their leaves and begin to emit other terpenes and additional compounds. This has been well characterised for cultivated *G. hirsutum*, but little is known about volatile production in response to herbivory in wild populations. We investigated how damage by the beet armyworm (*Spodoptera exigua*) affects leaf-stored and emitted volatiles in wild *G. hirsutum* plants. These plants were grown in a greenhouse using seeds collected from populations found along the Yucatan coast, from where this cotton species originates. We assessed whether the differences in leaf terpene profiles between two known chemotypes persisted upon herbivory, in leaves and in head-space emissions, and whether these chemotypes also differed in the production and release of herbivory-induced volatiles. In addition to chemotypic variation, we further investigated intraspecific variation in the volatile response to herbivory among genotypes, populations, and four geographic regions.

**Results:** The difference between the two chemotypes persisted after herbivory in the stored volatile profile of induced leaves, as well in the emissions from damaged plants. Therefore, wild cotton chemotypes may differ in their airborne interactions with their environment. The specific terpenes distinguishing these chemotypes showed a weak inducibility, raising questions about their functions. Herbivory triggered changes in stored and emitted volatiles similar to what is known for cultivated varieties of *G. hirsutum.* However, we report for the first time on the emission of volatile aldoximes by cotton plants, which were only detected in the headspace upon herbivory, and displayed chemotypic and interpopulation variation. Intraspecific variation was also observed in the induced emissions of nitriles and certain terpenes. Moreover, chemotypes differed in their induction of (*E*)-β-ocimene stored in the leaves.

**Conclusions:** This comprehensive insight into herbivore-induced volatiles of wild cotton reveals variation in production and emission among populations. A full understanding of their ecological role may help in the development of future pest-management strategies for cotton crops.

## Background

Plants emit volatiles that can play various defensive roles against insect herbivores. Volatile plant metabolites can function as toxins, repel insect feeding and oviposition, and attract natural enemies of their attackers [1,2]. Volatiles also mediate signaling within plants [3] and between neighboring plants, increasing their resistance against herbivores [4–6]. Unharmed plants can produce and emit certain volatiles constitutively, a phenomenon particularly prevalent in plant species that accumulate volatile terpenes in specialized structures. These stored terpenes are continuously emitted at low levels by intact plants and are released in much larger quantities when the storage organs are physically damaged, for example by chewing herbivores [7–9]. In addition, when plants detect an insect attack, they may actively synthesize socalled herbivore-induced plant volatiles (HIPVs), which can include compounds that are produced constitutively but synthesized in greater amounts upon damage, or compounds whose synthesis is triggered exclusively in response to herbivory [2,10].

Several wild plant species have been shown to exhibit intraspecific variation in their chemical phenotypes. This chemodiversity often has a genetic basis and can modulate the interactions between plants and their environment [11–13]. Intraspecific variation in volatile production has been described both for constitutive and herbivore-induced compounds [14,15]. Various species that constitutively produce and store volatile terpenes notably present discrete chemical polymorphisms in their terpene blend (chemotypes hereafter) [e.g. 14–18]. Importantly, this variation has been associated with differences in resistance to abiotic stress [21,22], allelopathy [23], defense against pathogens [24] and herbivores [24–28], as well as airborne signaling between plants when stored terpenes are released by damage [6,29]. Concerning HIPVs, variation has been reported for various classes of compounds across several wild species, such as: *Nicotiana attenuata* [30,31], *Datura wrightii* [32], *Solanum carolinense* [33,34], *Arabidopsis thaliana* [35,36], *Asclepias syriaca* [37], *Quercus robur* [38] and *Vicia sepium* [39]. Variation in HIPVs can be linked to differences in synthetic pathways [40,41], but also to differences in the endogenous signaling cascades that regulate the plants’ response to herbivory [31]. In addition, differences in the attraction of herbivores [34,38] and their natural enemies [34,36] have been found to be associated with variation in volatile induction.

Upland cotton (*Gossypium hirsutum* L. Malvaceae) is cultivated worldwide as a major source of natural fiber, and suffers from high levels of herbivory from multiple insect pests [42]. Increasing our understanding of the natural defense mechanisms of cotton could aid in the advancement of sustainable pest control strategies and the development of more resistant cultivars [2,43–45]. Cotton plants constitutively store volatile mono- and sesquiterpenes in specialized glands present in their leaf tissues [42,46]. Herbivory on older leaves induces an increase in the concentrations of these terpenes in young undamaged leaves [47,48]. These leaf-stored terpenes are steadily emitted at low levels from intact leaves, however large quantities are immediately released following gland rupture, for instance caused by chewing insects [49–52]. Therefore, volatile emissions instantly resulting from insect damage primarily comprise locally released leaf mono- and sesquiterpenes, together with green leaf volatiles (GLVs; aldehydes and alcohols), that are linked to tissue disruption [49,50]. Herbivory also induces a delayed emission of mono-, sesquiand homoterpenes, which are systematically emitted (i.e. also by undamaged parts) following their induced synthesis, and concurrently also triggers the emissions of other newly synthesized volatiles, such as the aromatic indole and GLV esters [49–55]. When active physical damaging ceases, the emission of constitutive leaf-stored terpenes drops rapidly, while induced compounds continue to be emitted [49]. The distinction between constitutive and truly inducible terpenes is not so clear cut, and some studies report systemic emissions of terpenes constitutively stored in leaves [54,55]. Collectively, volatile emissions of insect-damaged cotton plants appear to serve as direct and indirect defenses [reviewed in 42], and are involved in plant-plant interactions .

Domestication often leads to a reduction in defenses against herbivory in crop plants [59,60]. Modern cultivars of *G. hirsutum* exhibit a depletion of genetic diversity [61] and also appear to be less resistant to herbivory compared to wild-growing plants [62–65]. Therefore, exploring useful defensive traits in wild populations of *G. hirsutum* should be considered. While volatile production in cultivated varieties has been well characterized, less is known about the volatiles produced by wild *G. hirsutum* plants in response to herbivory, and studies conducted so far mostly relate to naturalized varieties [62,63,66]. *G. hirsutum* was domesticated at least 4,000 years ago in the Yucatán Peninsula (southeast Mexico) [61], where wild populations still grow naturally along its northern coastline [67,68].

Recently, we described two heritable constitutive leaf-terpene chemotypes in wild cotton plants from the Yucatan coast: one chemotype containing either none or low amounts of γ-terpinene (and other co-expressed monoterpenes), and the other chemotype exhibiting distinctively higher abundances of these compounds in their leaves [69]. It is worthwhile to note that production of γ-terpinene has been only described in wild/naturalized cotton and from crosses between cultivated and wild cotton [62,63,66]. If γterpinene variation in leaves influences the composition of the volatile emissions of wild cotton plants, it could impact the plants’ interactions with organisms that use these volatiles as a cue, such as herbivores looking for a host plant, their natural enemies, or neighboring plants.

The present study investigated the effect of damage by caterpillars (beet armyworm, *Spodoptera exigua* Hübner; Lepidoptera; Noctuidae) on both the volatile terpenes stored in leaves and the volatile emissions of plants from wild populations of *G. hirsutum*. Plants were grown in a greenhouse from seeds collected from sixteen populations forming four distinct geographic zones distributed along the Yucatan coastline. We assessed the consistency of the two leaf-terpene chemotypes, previously described as constitutive, in the stored volatile profile of induced leaves, as well in the emissions of both undamaged and damaged plants. In addition, we evaluated the variations in the induction by herbivory of other stored and released volatiles between the two chemotypes, and among three nested levels of intraspecific variation: genotypes, populations, and the geographic regions grouping the populations. As the prevalence of the two chemotypes along the coast follows a geographic gradient [69], chemotypic variation is interrelated with variation among populations.

The results presented here shed new light on the production of volatiles by wild *G. hirsutum* plants in response to caterpillar herbivory, and uncover sources of variation in volatile induction, which may be genetically based, thereby opening new avenues to study and exploit cotton HIPVs as a defense mechanism against insect pests.

## Methods

### 1 Plants and insects

Wild *Gossypium hirsutum* seeds were collected in 2019 and 2020 on the northern coast of the Yucatán Peninsula (Mexico) and grown as described in [69]. From July to September, plants were grown in individual pots filled with 250 ml of soil (Profi Substrat, Einheitserde, Germany), randomly arranged within a single greenhouse located in Neuchâtel. Seeds were sampled from several mother plants from 16 populations in four regions along the coast (Table S1 and Fig. 1). As cotton cross-pollinates, seeds from the same mother are either full or half-siblings, of unknown paternal source. Offspring from the same mother plant is referred to here as genotype. Plants were grown in a greenhouse for approximately four weeks. Eventually, we obtained plants from one to six mother plants for each population, representing around 35 to 60 individuals per population, except for a few populations with particularly low germination rates (Table S1). To induce volatile production in the cotton plants, we used larvae of the moth *Spodoptera exigua* (Hübner; Lepidoptera; Noctuidae), a cosmopolitan generalist pest that also feeds on cotton. Eggs were supplied by Entocare (Wageningen, The Netherlands) and reared in the laboratory on wheat-germ based artificial diet (Frontier Scientific Services, Newark, USA).

**Figure 1.**
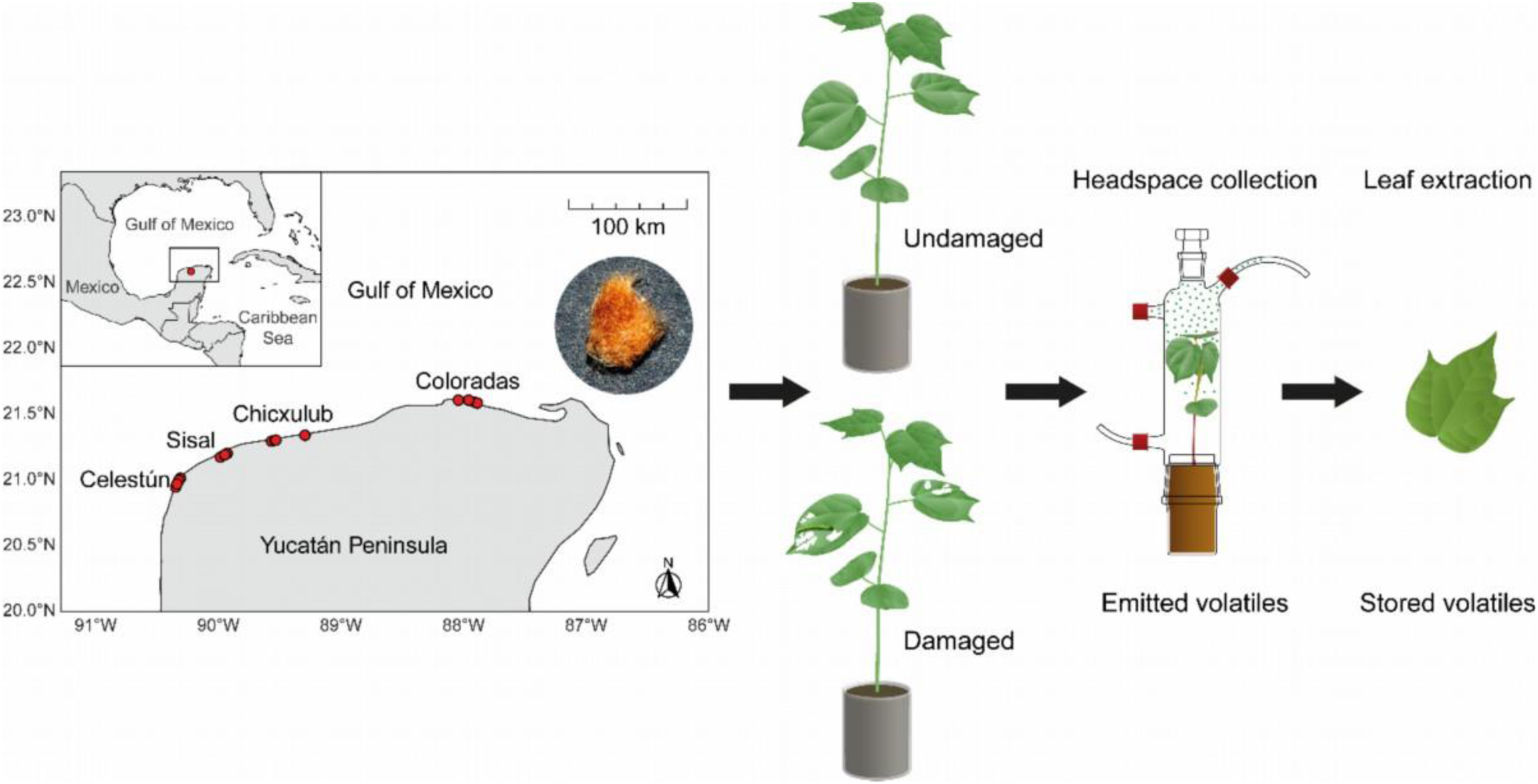
Illustration of the experimental process. Seeds were collected from 16 populations (red points) in four regions along the northern coast of the Yucatán Peninsula. Plants were grown in a greenhouse until they developed the fourth leaf, at which time half of the plants were damaged by *Spodoptera exigua* larvae on their first and second leaves for two days and the other half was kept intact. Emitted volatiles were collected by headspace sampling and stored leaf volatiles by hexane extraction of the undamaged fourth leaves.

### 2 Experimental process and sample collection

Plants were used when the fourth leaf was fully expanded but still developing. A total of 322 plants were subjected to herbivory and 353 were left undamaged, in six experimental batches. The numbers of replicates per region and population are described in Table S1. Second to early third-instar *S. exigua* larvae fed on the first and second leaves for two days, with a single larva enclosed on each leaf in clip cages (4.52 cm^2^, BioQuip product, USA). Undamaged and damaged plants were kept in two adjacent rooms with similar configurations and conditions (L14:D10, T=28°C), to prevent any signaling effect of HIPVs. The treatments were alternated between rooms after each batch. After two days of herbivory, we collected the volatile emissions of a subset of the plants. A total of 45 undamaged and 48 damaged individuals were used in four batches for headspace collection, chosen to represent the various populations (Table S1). Afterwards, the fourth leaf of all plants was harvested, photographed, frozen in liquid nitrogen, and stored at -80 °C until processed. Photographs of all leaves were taken to quantify the total leaf area consumed by *S. exigua*, as well as to determine the area (size) of the fourth leaves and the total leaf area (including cotyledons and secondary leaves) of the plants used for the headspace collections. For this purpose, we used the Fiji distribution of ImageJ software [70].

### 3 Headspace volatile collection and GC-MS analyses

*S. exigua* larvae were removed before headspace sampling, to avoid the release of volatiles due to physical tissue disruption by active feeding. The caterpillars’ frass was cleared away from the leaves. Pots were covered with aluminum foil to limit potential emissions from the soil. Headspace volatiles were collected in a push-pull air system of custom-made glass bottles (VTQ SA, Switzerland). Charcoal-purified and humidified air was pushed at 1 L min^-1^ and pulled at 0.6 L min^-1^ through trapping filters (25 mg of 80/100 mesh Hayesep-Q adsorbent, Sigma, Switzerland) for two hours. Filters were eluted with 100 μL of dichloromethane, with the addition of 10 μL of internal standard solution (nonyl-acetate, 20 ng μL^-1^). Samples were stored at -80 °C before being run on a gas chromatograph (GC Agilent 7890B) coupled to a mass spectrometer (MSD Agilent 5977B): 1.5 μL were autoinjected in splitless mode onto an Agilent HP-5MS column (30 m × 250 µm × 0.25 µm). The temperature was kept at 40 °C for 3.5 min, then ramped to 100 °C at 8 °C min^-1^ and to 230 °C at 5 °C min^-1^. Helium was used as the carrier gas, with a stable flow of 0.9 mL min^-1^.

### 4 Hexane leaf extractions and GC-MS analyses

Hexane leaf extractions and subsequent analyses were performed as described in Clancy et al. (2023). Briefly, 100 mg of finely ground frozen leaf material was extracted with 1 mL of hexane and kept at 4 °C for 24 hours. 800 μL was recovered and 8 μL of internal standard solution (nonyl acetate, 2000 ng μL^-1^) was added. The samples were directly injected into a gas chromatograph (GC Agilent 8890A) coupled to a mass spectrometer (MSD Agilent 5977B): 1.5 μL were injected in splitless mode onto an Agilent HP-5MS column (30 m × 250 µm × 0.25 µm). Temperature started at 40°C for 0 min, ramped to 130 °C at 15 °C min^-1^, then to 190 °C at 60 °C min^-1^, and to 290 °C at 30 °C min^-1^, then held for 0.5 min. Helium was used as carrier gas, with a stable flow of 0.9 mL min^-1.^

### 5 GC-MS data processing

PyMS (O’Callaghan et al., 2012) was used to integrate and align peaks from the total ion chromatograms of all samples (https://github.com/bustossc/PyMS-routine/). Analytes were quantified using relative response factors obtained from a series of external calibration curves of commercial analytical standards in comparison to nonyl acetate (Table S2 and S3). For compounds lacking available response factors, we employed the relative response factor associated with the closest standard in retention time and similar chemical structure, while normalizing by molecular masses (Kreuzwieser et al. 2014). When possible, analytes were identified based on comparison with commercial standards. Aldoximes were identified with synthesized standards kindly provided by Dr. Tobias Köllner. Compounds with no standard available were identified based on mass spectra comparison with NIST 2.3 mass spectral library, an in-house library, and Kovats retention indices [71] (Table S2 and S3). Components retained in the analyses were present in at least 10% of the samples. For the hexane extracts, values were normalized by the fresh weights of the leaves (units: ng mg^-1^ FW). For headspace samples, values were normalized by hours of collection and the total leaf area of the plants (units: ng h^-1^ cm^-2^).

### Statistical analyses

Statistical analyses were carried out with R 4.3.2 [72]. A significance level of α = 0.05 was used for all analyses. Plots were built with “ggplot2” [73] (v. 3.4.4).

#### 1 Multivariate analyses of volatiles composition: pRDAs and correlation analyses

Multivariate analyses were always performed on the proportions of compounds relative to the sum of all analytes of interest. To deal with the left-censored and compositional nature of the data, zeroes were imputed by multiplicative simple replacement with multRepl() from “zCompositions” [74] (v. 1.5.0.1), using the smallest non-zero value of each compound as detection limits and a delta of 0.65. We subsequently employed a centered log-ratio (clr) transformation with clr() from “compositions”[75] (v. 2.0.8). Data were finally scaled and centered using scale() from the R base package. To evaluate correlations among compounds, hierarchical clustering was performed on the correlation-based (Pearson) dissimilarity matrix with Ward’s minimum variance method (Ward.D2) using pvclust() from “pvclust” [76] (v. 2.2.0). Approximately Unbiased (AU) p-values were computed by multiscale bootstrap resampling with 10,000 replications. Correlations heatmaps were plotted with “ComplexHeatmap” [77] (v. 2.16.0). Partial redundancy analyses (pRDAs) were conducted with rda() from “vegan”[78] (v. 2.6.4). Models’ structures are described in the following paragraphs. We then performed a type II permutation test with MVA.anova() from “RVAideMemoire”[79] (v. 0.9.83.7). Pairwise comparisons were performed with pairwise.factorfit() from the same package.

#### 2 Analyses of single compounds: GLMMs

Univariate analyses of concentrations of compounds in leaves or emission rates were performed with generalized linear mixed models (GLMMs) using glmmTMB [80](v. 1.1.9) with Gamma or Tweedie exponential families of distribution, depending on models’ fit. Models’ fits were checked with the package “DHARMa”[81] (v. 0.4.6). Effects of the explanatory variables were tested with Wald chi-squared tests (type-II) using Anova() from “car” [82] (v. 3.1.2). We plotted estimated marginal means and their 95% confidence intervals (CI), extracted from the models with ggemmeans() from “ggeffects” [83] (v. 1.3.4). When plotted, data points represented observed values. In all models, we controlled for the effects of plants’ “size” (leaf extracts: fourth leaf area; headspace emissions: total leaf area) and of experimental batches (leaf extracts: n = 6; headspace emissions: n = 4) by including them as fixed effects (Table S2 and S3). γ-Terpinene and coexpressed compounds were analyzed within the producing chemotype only, as the other chemotype either does not produce them, or only in very low amounts. Population and genotype were included as random intercepts in all models, except for those relying solely on the headspace emissions of damaged plants, where 86% of the data consisted of individual data points per genotype (Table S2 and S3). When necessary, pairwise comparisons were performed with emmeans() from “emmeans”, with multiple test adjustment (FDR)[84] (v. 1.10.0).

#### 3 Volatile terpenes in leaves: herbivory and variation in induction

To assess how herbivory affected the leaf composition in volatile terpenes for the two chemotypes, a pRDA was performed on the relative abundances of all leaf mono- and sesquiterpenes. Herbivory, chemotype and their interaction were used as explanatory variables while leaf area was partialled out. Then, we analyzed the effect of herbivory and damage intensity (total leaf area consumed by caterpillars) on the total concentrations of the leaf volatile terpenes and a subset of compounds using GLMMs (Table S2: model structures A and E). We tested whether there was variation in the induction (i.e. increase in the leaf concentrations; group-by-herbivory interaction) of volatile terpenes between the two chemotypes and among nested levels of intraspecific variation: 88 genotypes, 16 populations and four geographic regions. Variation in the response of genotypes and populations to herbivory was modelled with random slopes and their significance was tested by comparing nested models with a random slope component to a model without (Restricted Maximum Likelihood method) with Likelihood ratio tests. Chemotype was included in the fixed effects of these models (Table S2: model structures B and C). The random slope component included a term for the variance in slopes and one for the covariance between slopes and intercepts unless the model did not converge. In that case, the covariance term was removed. Variation in response to herbivory among region and chemotypes was modelled as interaction between these groups and herbivory, set as fixed effects). Because of their geographic distribution, chemotypes were partially nested in regions. Therefore, to avoid collinearity, they were always analyzed separately (Table S2: model structures A and D).

#### 4 Headspace volatile emissions: herbivory and variation in induction

To examine whether the intraspecific variation observed in the leaf-stored terpenes by Clancy et al. (2023) was reflected in the emissions, we performed Spearman correlation tests, separately for undamaged and damaged plants, on the ratio of concentrations of γ-terpinene and α-pinene (the dominant leaf-stored monoterpene) in leaves and in headspace emissions. To visualize the composition of the emitted blends of undamaged and damaged plants of both chemotypes, we plotted the median proportions with “treemapify” [85] (v. 2.5.6). To further understand how herbivory and chemotypes affected the composition of headspace emissions a pRDA was conducted on the relative abundances of all emitted volatiles together. Herbivory, chemotype and their interaction were used as explanatory variables while the total leaf area was partialled out. Then, we analyzed the effect of herbivory and damage intensity (total leaf area consumed by caterpillars) on the total emission rates and on the emissions of same terpenes analyzed for the leaves (Table S3: model structures A and C).

Variation in the volatile emissions induced by caterpillar herbivory was evaluated between chemotypes, as well as two levels of intraspecific variation: among the four regions and 16 populations. Genotypes were not assessed because of the lack of replication at this level. To evaluate variation in the volatile response to herbivory rather than pre-existing differences in compounds already emitted by intact plants, we selected only highly inducible volatiles, defined from the pRDA assessing the effect of herbivory on the headspace emissions. We first evaluated whether there was variation in the total emissions of these highly inducible compounds (Table S3, model structures C, D and E). Variation among population was modelled through random intercepts (Table S3: model structures E) and their significance was tested by comparing nested models with random intercept component to models without (Restricted Maximum Likelihood method) with Likelihood ratio tests. Chemotypes and regions were analyzed as fixed effects, and separately, as mentioned above. To unveil potential variation in the composition of the induced emissions, pRDAs were performed on the proportions of highly inducible volatiles, selected from the transformed and scaled matrix of the relative abundances of all compounds emitted by damaged plants, separately for chemotype and region. Chemotype/region, the leaf surface damaged and their interaction were used as explanatory variables while the total leaf area was partialled out. We then further tested whether chemotypes and regions varied significantly in their emission rates of the different candidate compounds highlighted by the pRDAs with GLMMs, as explained for the total induced emissions (Table S3, model structure C and D). Compounds with significant variations were also assessed for variation among populations (Table 4, model structure E). For inducible compounds differing between chemotypes or regions but also emitted by undamaged plants, the interaction effect between chemotype/region and herbivory was tested (Table S3, model structure A and B). A significant interaction implied that chemotypes/regions differed in their response to herbivory, not solely by pre-existing disparities.

## Results

### 1 Leaf-stored volatiles

We extracted volatile content from the leaves of undamaged (n=353) and damaged (n=322) wild cotton plants (Fig. 1). A total of 15 monoterpenes, 18 sesquiterpenes and three GLVs were quantified in the leaf extracts (Table S4). Subsequent analyses focus on terpenes.

#### 1.1 Monoterpene chemotypes

Chemotypes were assigned to each plant based on the leaf volatile profile. First, undamaged plants were chemotyped based on the contribution of six strongly correlated compounds relative to the total monoterpene concentration (γ-terpinene, limonene, α-thujene, terpinolene, p-cymene, and α-terpinene, in order of abundance; *sensu* [69]; Fig. S1, Table S4). Plants containing higher proportions (> 4%) of these compounds were assigned to chemotype A; plants with less than 4% were assigned to chemotype B (Fig. 2). The six variable monoterpenes remained highly correlated upon the two days of herbivory (Fig. S2) and damaged plants could be separated in two chemotypes in the same way as undamaged plants (Fig. 2).

**Figure 2.**
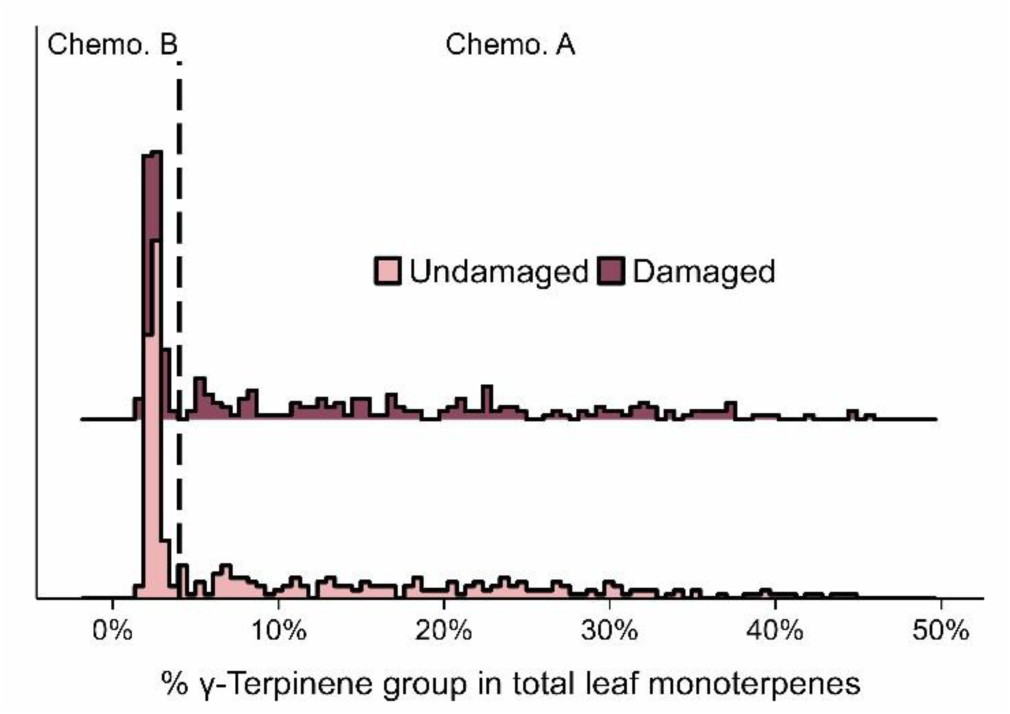
Chemotypic variation associated with γ-terpinene in leaf volatile profiles of wild cotton plants. Proportion of γ-terpinene, limonene, α-thujene, terpinolene, p-cymene, and α-terpinene («γ-terpinene group») among monoterpenes stored in wild cotton leaves of undamaged and damaged wild cotton plants.

#### 1.2 Effect of herbivory on the leaf-stored volatile terpene profiles

Following herbivory during two days on the first and second leaves of four-leaf plants, the total leaf concentration of mono- and sesquiterpenes increased by 9% on average in the undamaged fourth leaves (Fig. 3A, χ^2^(1) = 12.407, p < 0.001). The same compounds were detected in the leaf terpene profiles of undamaged and damaged plants (Table S4), but a partial redundancy analysis (pRDA) revealed that herbivory significantly affected their relative abundances (Fig. 3B, F (1,670) = 6.414, p = 0.001). This change in composition upon herbivory appeared to be mainly linked to the increase in the proportion of the monoterpene (*E*)-β-ocimene, followed by several sesquiterpenes, notably δ-cadinene and α-copaene (Fig. 3B). The induction of (*E*)*-*β-ocimene depended on chemotype (Fig. 3A; chemotype x herbivory: χ^2^(1) = 7.694, p = 0.005), with a 64% increase in chemotype A plants (contrast of EMMs: p < 0.001) and a 108% increase in chemotype B plants (contrast of EMMs: p < 0.001). Leaf content of the sesquiterpene δ-cadinene increased following herbivory by 27% (Fig. 3A, χ^2^(1) = 27.201, p < 0.001). (*E*)*-*β-caryophyllene, the most abundant sesquiterpene in leaves (Table S4), increased by 11% (Fig. 2A: χ^2^(1) = 14.254, p < 0.001). In contrast, levels of α-pinene, the most abundant monoterpene in leaves (Table S4), did not change upon herbivory (Fig. 3A, χ^2^(1) = 0.437, p = 0.509). This was also the case for γ-terpinene, the main monoterpene characteristic for chemotype A (Fig. 3A, χ^2^(1) = 0.766, p = 0.381). The total accumulation of (*E*)*-*β-ocimene increased with the amount of leaf tissue consumed by *S. exigua,* but this effect of damage intensity was not significant for the other terpenes tested (Fig. S3: (*E*)*-*β-ocimene: χ^2^ = 5.619, p = 0.018).

**Figure 3.**
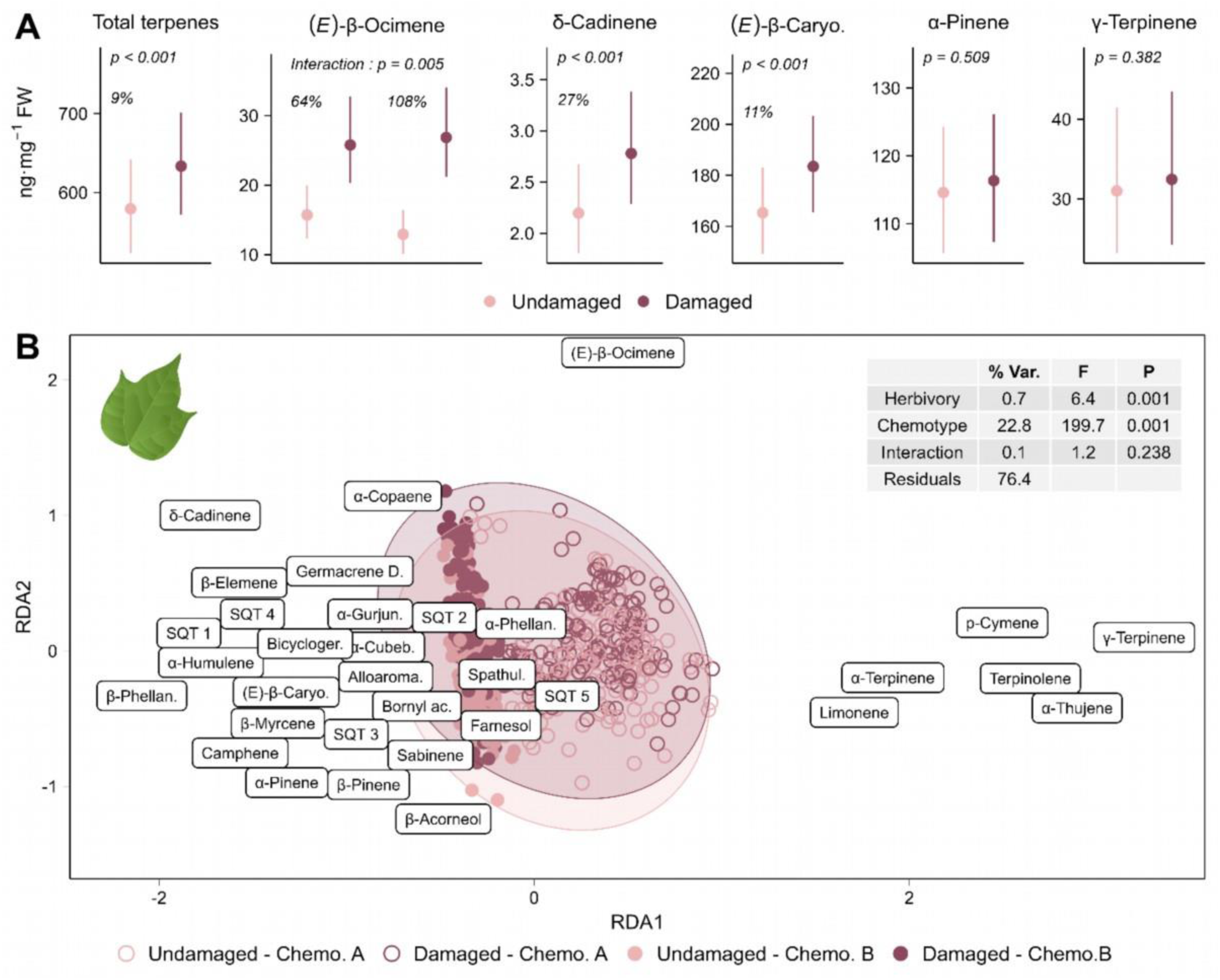
Effect of herbivory on stored leaf monoand sesquiterpenes for both wild cotton chemotypes. Volatile compounds were extracted from the fourth leaves of 353 undamaged wild cotton plants and 322 plants damaged by *Spodoptera exigua* for two days on their first and second leaves. **A:** Effect of herbivory on the leaf concentrations of a subset of terpenes. Estimated marginal means and 95% CI from GLMMs. γTerpinene was analyzed within chemotype A while all other compounds were analyzed for both chemotypes together, with chemotype set as fixed factor. **B**: Biplot with symmetric scaling of a pRDA on the relative abundances (clr-transformed) of leaf monoand sesquiterpenes. It elucidates the relationship between volatile composition and herbivory, chemotype and their interaction, while accounting for the effect of leaf area. The proximity of the points indicates the similarity in volatile composition. The table provides a summary of the permutation test results, with the percentage of variation explained by each variable (% Var.), the F-statistic (F) and the p-value (P).

#### 1.3 Variation in the induction of leaf-stored volatiles

We investigated whether there was intraspecific variation in the volatile response to herbivory for the total leaf terpene content and for the aforementioned compounds. As volatile leaf terpenes are constitutively produced, we assessed the variation in the increase of leaf concentrations induced by herbivory, i.e. a group-by-herbivory interaction, groups being either chemotypes, genotypes, populations or regions (Table S5). As mentioned above, induction of (*E*)*-*β-ocimene varied between the two chemotypes (Fig. 3A and Table S5; chemotype x herbivory: χ^2^ = 7.694, p = 0.005), with a stronger induction observed in chemotype B plants. Chemotype A plants exhibited constitutively higher amounts of (*E*)*-*β-ocimene in their leaves than chemotype B plants (Fig. S4, χ^2^ = 8.247, p = 0.004); this difference was no longer significant following herbivory (Fig. S4, χ^2^ = 0.793, p = 0.373). We did not find any significant variation in induction of any tested compounds among genotypes, populations, and regions (Table S5).

### 2 Headspace volatile emissions

The volatile emissions of undamaged and damaged wild cotton plants were collected and characterized in a subset of the plants that were afterwards used for leaf extraction (Fig. 1; 45 undamaged and 48 damaged plants). We quantified a total of 54 compounds in the headspace (Table S6). Terpenes represented the richest group, with 13 monoterpenes, 13 sesquiterpenes, and the homoterpenes DMNT and TMTT identified. Additionally, headspace emissions contained many fatty acids derivatives, which comprised GLV aldehydes, alcohols, and esters, and (*Z*)-jasmone (Table S6). Also, we identified three aliphatic aldoximes and several aromatic compounds including two nitriles. Most of the mono- and sesquiterpenes detected in the headspace were also found in the leaf extracts, as well as a GLV aldehyde (Fig. 4A).

**Figure 4.**
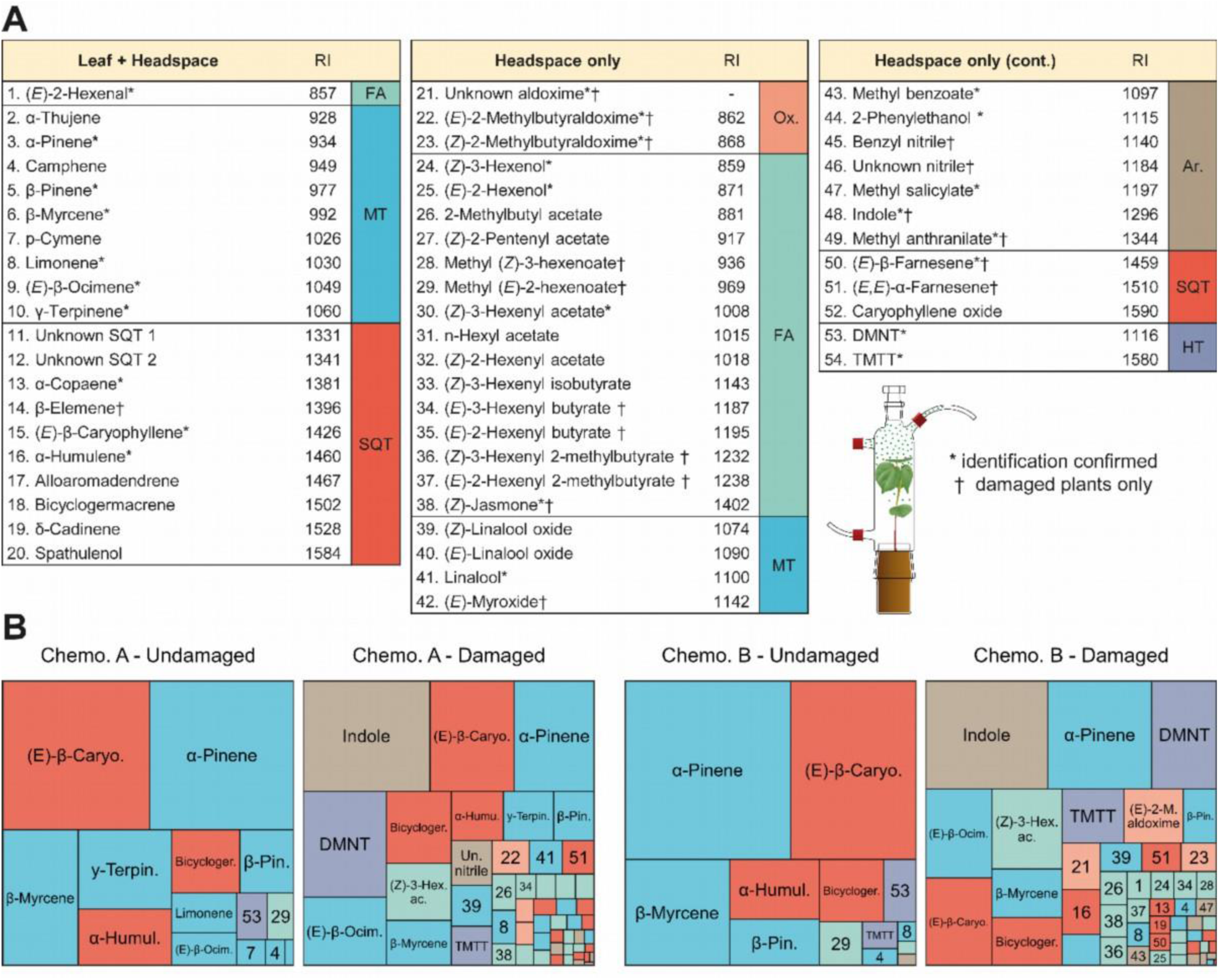
**Relative composition of the headspace volatiles emitted by undamaged and damaged wild cotton plants**. **A:** Compounds detected both in the plants’ leaves extracts and headspace, or in their headspace only. RI = Retention indices. The retention index for the unknown aldoxime could not be calculated because it eluted before the smallest alkane. † indicates compounds specific to damaged plants (detected in less than 10% of undamaged plants, but in between 43% and 100% of damaged plants). * indicates compounds whose identity was confirmed with standards. Distinction between the two 2-methylbutyraldoximes isomers is tentative. FA = fatty acid derivatives; MT = monoterpenes; SQT= sesquiterpenes; Ox. = aldoximes; Ar. = aromatics; HT = homoterpenes. **B:** Composition of medians emitted volatile profiles of undamaged and damaged plants of both chemotypes. Area of the rectangles is proportional to the contribution of the compound to the profile. Numbers refer to A.

#### 2.1 Monoterpene chemotypes

We assessed whether the leaf-terpene chemotypes could also be distinguished from their headspace emissions, both for undamaged and damaged plants. Of the six monoterpenes that were used to define chemotypes A and B using leaf extracts, the following compounds were also detected in the headspace emissions (in order of abundance): γ-terpinene, limonene, p-cymene, and α-thujene (Table S6). For both undamaged and damaged plants, the ratio of γ-terpinene to α-pinene in headspace emissions was correlated to their ratios in the leaves (Fig. 5) and volatile emissions of the two chemotypes were clearly distinguishable by the prevalence of γ-terpinene, p-cymene, and α-thujene (Fig. 4B and 6A), which were highly correlated (Fig. S5 and S6).

**Figure 5.**
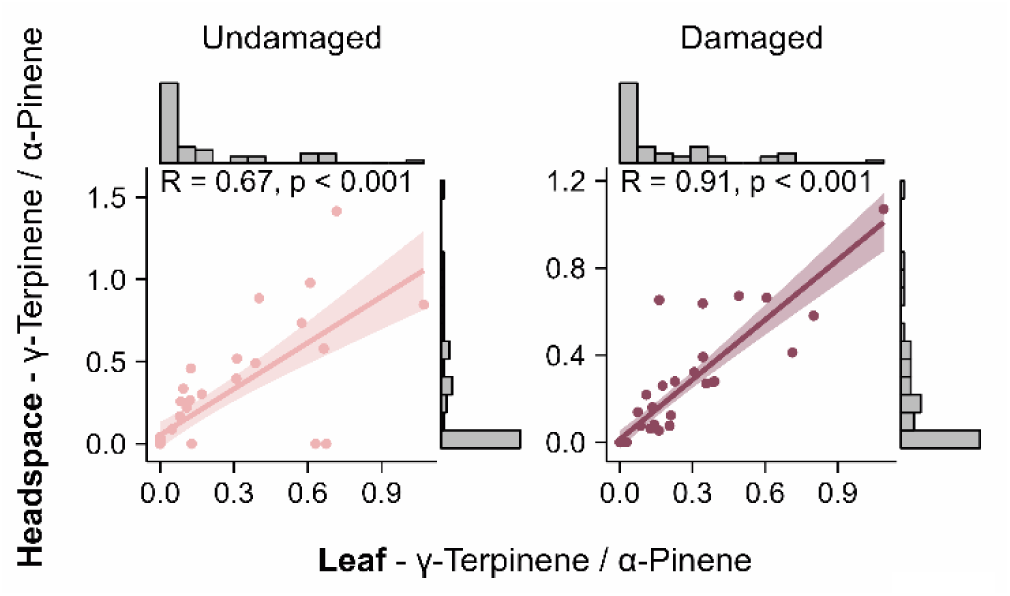
Effect of γ-terpinene variation in leaves on headspace emissions: correlation (Spearman) between the ratio of γ-terpinene and α-pinene concentrations in leaves (x-axis) and in headspace emissions (y-axis).

**Figure 6.**
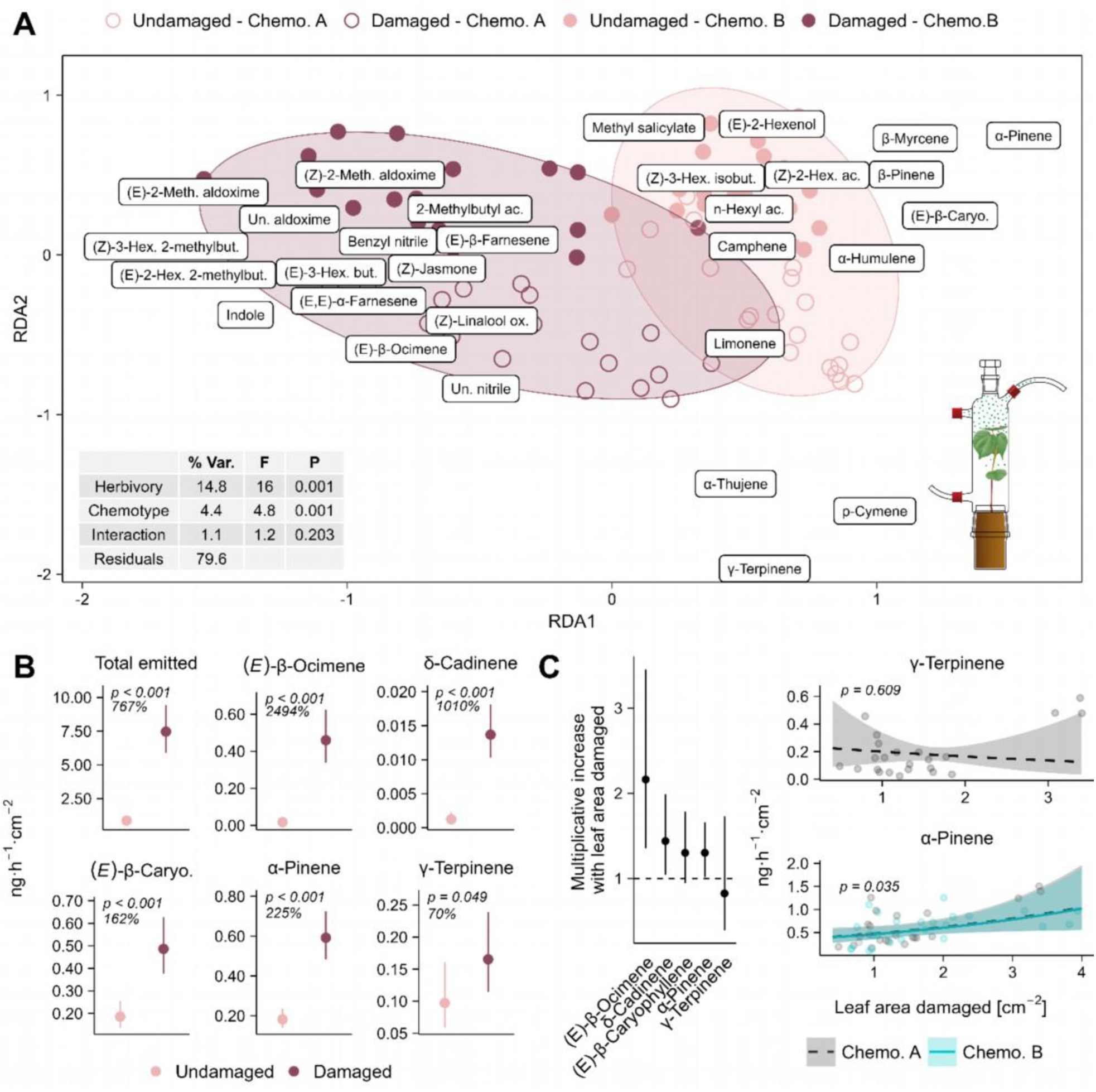
Effect of herbivory on the volatiles emitted by the two wild cotton chemotypes. Headspace volatiles were sampled from 45 undamaged wild cotton plants and 48 plants damaged by *Spodoptera exigua* for two days on their first and second leaves. **A:** Biplot with symmetric scaling of a pRDA on the relative abundances (clr-transformed) of all compounds quantified in the headspace samples. Only compounds accounting for more than 10% of inertia are plotted (full biplot in Fig. S7). The plot elucidates the relationship between volatile composition and herbivory, chemotype and their interaction, while accounting for the effect of the total leaf area of the plants. The proximity of the points indicates the similarity in volatile composition. The table provides a summary of the permutation test results, with the percentage of variation explained by each variable (% Var.), the F-statistic (F) and the p-value (P). B: Effect of herbivory on emission rates of all emitted volatiles and of a subset of emitted terpenes found also in the leaf extracts. Estimated marginal means and 95% CI from GLMMs are plotted. γ-Terpinene was analyzed within chemotype A while all other compounds were analyzed for both chemotypes together, with chemotype set as fixed factor. C: relationship between damage intensity (leaf area damaged) and the emission rates of a subset of terpenes. Left: multiplicative increases with the leaf area damaged (backtransformed coefficient estimates from GLMMs and their 95% CIs). The dotted line indicates a coefficient of 1, where quantities emitted remain constant with damaged area. Right: adjusted predictions and 95% CI from GLMMs with observed values as data points. γ-Terpinene was analyzed within chemotype A while all other compounds were analyzed for both chemotypes together, with chemotype set as fixed factor.

#### 2.2 Effect of herbivory on the emitted volatile profile

The volatile blend emitted by undamaged plants was mainly composed of mono- and sesquiterpenes that were also found in leaf extracts, with (*E*)*-*β-caryophyllene and α-pinene dominating the median profile of both chemotype A and B plants (Fig. 4B). Herbivory caused the total volatile emissions to increase 8.7-fold (Fig. 6B, χ^2^ (1) = 146.449, p < 0.001) and triggered qualitative and quantitative changes in their composition (Fig. 4, 6A and S7). Damaged plants released several compounds that were absent from the headspace of intact plants, specifically aliphatic aldoximes, benzyl nitrile and another unidentified nitrile, the aromatics indole and methyl anthranilate, butyric and hexenoic acid esters of GLVs, (*Z*)-jasmone, the acyclic sesquiterpenes (*E*)*-*β-farnesene and (*E,E*)*-*α-farnesene, and monoterpene (*E*)-myroxide (Fig. 4). Other compounds such as the acyclic monoterpenes (*E*)*-*β-ocimene, linalool and (*Z*)-linalool oxide, the homoterpenes DMNT and TMTT, and the GLV ester (*Z*)*-*3-hexenyl acetate were emitted in low amounts by some of the undamaged plants, but their prevalence greatly increased following herbivory (Fig. 4B, 6A and S7). Indole, DMNT, (*Z*)*-*3-hexenyl acetate, and (*E*)*-*β-ocimene were major constituents of the volatile blend from damaged plants of both chemotypes (Fig. 4B). The emission rates of the mono- and sesquiterpenes that were also found in the leaf extracts generally exhibited small increases upon herbivory (Fig. 6B). However, due to greater increases of other compounds, their relative contribution to the emissions of damaged plants decreased (Fig. 4B and 6A). This was not the case of (*E*)*-*β-ocimene, which exhibited a distinctly high increase in response to herbivory compared to the other terpenes found in both headspace and leaves. Irrespective of chemotype (χ^2^(1) = 1.901, p = 0.168), the amounts emitted of (*E*)*-*β-ocimene rose 26-fold in response to herbivory (Fig. 6B, χ^2^ (1) = 139.057, p < 0.001). In contrast, (*E*)*-*β-caryophyllene and α-pinene only increased 2.6-fold (Fig. 6B, χ^2^(1) = 33.800, p < 0.001) and 3.2-fold (Fig. 6B, χ^2^(1) = 56.859, p < 0.001) respectively. Emissions of δ-cadinene showed an intermediate increase of 11-fold (Fig. 6B, χ^2^ (1) = 62.593, p < 0.001). A particularly low response was observed for γ-terpinene, with only a marginal increase in the headspace of 1.7-fold following herbivory (Fig. 6B, χ2(1) = 3.868, p = 0.049).Emissions of γ-terpinene by damaged plants were independent of damage level (Fig 6C, χ^2^(1) = 1.299, p = 0.254), as with p-cymene, α-thujene, and limonene (Fig. S8). The emissions of all other terpenes detected in both headspace and leaf extracts were positively correlated with damage intensity (Fig. S8).

#### 2.3 Variation in the induction of volatile emissions

We investigated whether there were differences in the volatile emissions induced by *S. exigua* between chemotypes, as well as among regions and populations. Genotypes were not assessed because of the low level of replication. Our analyses focused on highly inducible volatiles. These were defined as the compounds with a negative score on the first axis, which separated damaged and undamaged plants, of the pRDA in Figure 5A (see Fig. S8 for a biplot with all compounds displayed). Total emissions of these highly inducible compounds did not differ between chemotypes (Fig. S9, χ^2^(2) = 0.003, p = 0.954), nor among regions (Fig. S9, χ^2^(3) = 2.893, p = 0.408) or populations (Fig. S9, χ^2^(1) = 1.344, p = 0.246). To further investigate differences between chemotypes and among regions, a new set of pRDAs were conducted on the abundances of these highly inducible compounds relative to the whole volatile blend emitted by damaged plants. These analyses revealed differences between chemotypes (Fig. S10, F (1,42) = 1.189, p = 0.009) and among regions (Fig. S11, F (3,38) = 1.904, p = 0.005) in the composition of their HIPV emissions. Of the compounds highlighted by the pRDAs, the terpenes (*E,E*)-α-farnesene, (*E*)-myroxide, the three aldoximes and the two nitriles showed statistically significant differences in emission rates between chemotypes and/or regions (Table S7). These compounds were all only detected in the headspace of damaged plants. We also observed variations in the emissions of TMTT and δ-cadinene (Fig. S10 and S11); these volatiles were also found in undamaged plants but in lower amounts. The absence of significant interactions between region/chemotype and herbivory suggests that the observed differences might not be specifically linked to a distinct response to herbivory (Fig. S10 and S11). Emission rates of (*E,E*)-α-farnesene and (*E*)-myroxide varied among regions, as well as among populations, but not between chemotypes (Table S7, Fig. 7 and Fig. S9). The three aldoximes, which were strongly correlated (Fig. S6), varied both among regions and between chemotypes. Plants from Celestún (far west) and Sisal (west) produced higher quantities of aldoximes compared to plants from regions further east (Table S7, Fig. 6 and Fig. S9). Chemotype B plants released higher amounts of these compounds than chemotype A plants (Table S7, Fig. 6 and Fig. S9). Thus, variation could stem both from the chemotype and the geographic region, which are partially nested. However, even when considering the effect of chemotype in the models, significant variation was observed at the population level for the two methylbutyraldoximes (Table S7). Benzyl nitrile showed the same trend in variation as the aldoximes, while the opposite was observed for the unknown nitrile (Table S7, Fig. 6). For these latter two compounds, however, no evidence of variation among populations was observed (Table S7).

**Figure 7.**
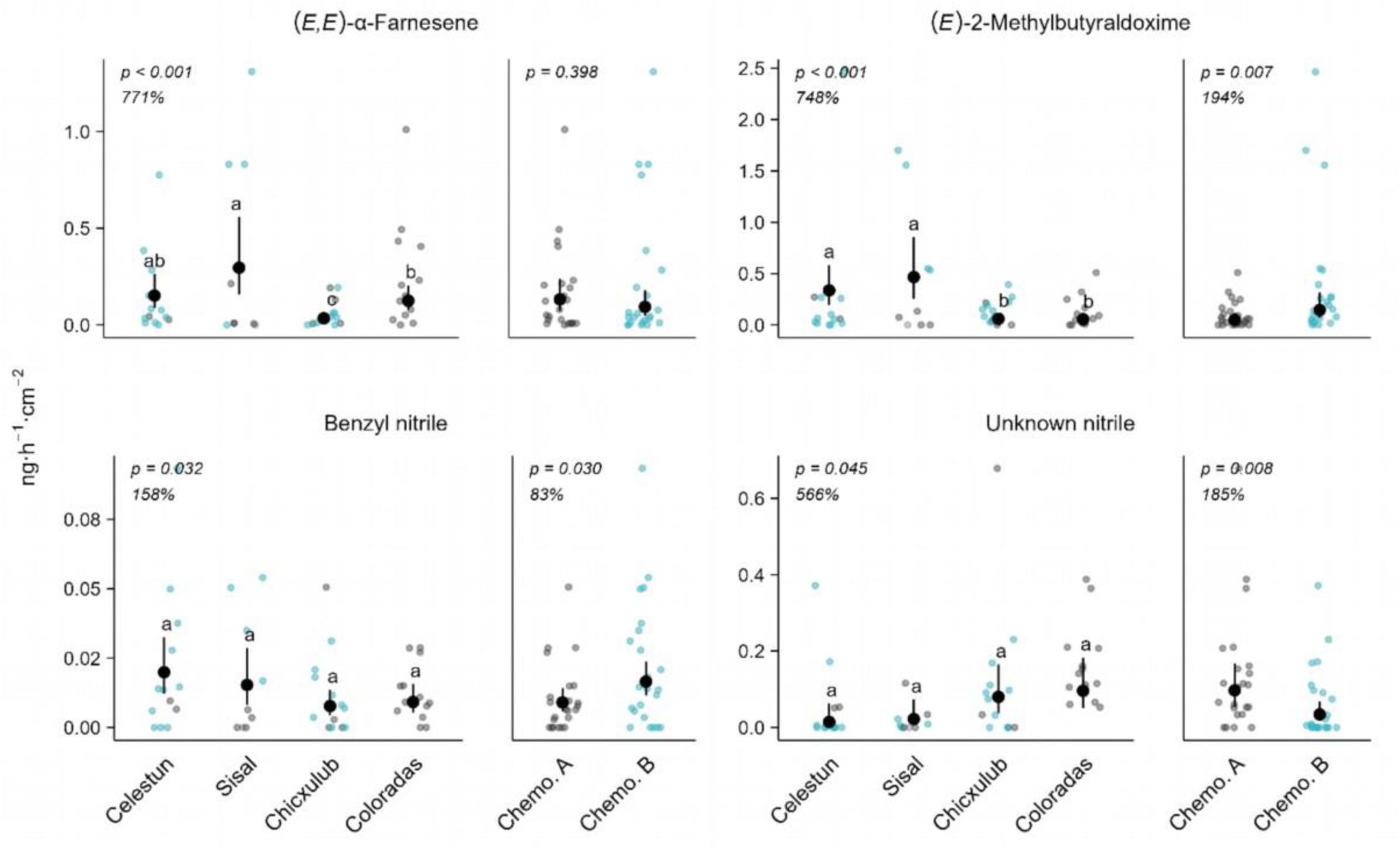
Variation in emitted HIPVs among geographic regions and between wild cotton chemotypes. Headspace volatile emissions were sampled from 48 wild cotton plant damaged by *Spodoptera exigua* for two days. Regions are displayed from west to east. Prevalence of chemotypes is nested with regions, as indicated by the color of data points (grey points indicate chemotype A; blue points indicate chemotype B), therefore they were analyzed separately. Estimated marginal means and 95% CI from GLMMs with observed values as data points. Damage intensity was controlled for as covariate. Letters indicate significant pairwise comparisons after multiple test adjustment (FDR). For regions, % values indicate the difference between the region with the lowest and highest estimated mean emission rates.

## Discussion

### 1 Consistency of the leaf-terpenes chemotypes

The two previously reported constitutive leaf-terpene chemotypes [69] were also distinct in their leaf volatile terpene profile after herbivory. They were also distinctive in their headspace emissions, both from undamaged and damaged plants. The fact that the variation in the leaf content of γ-terpinene and its correlated compounds was mirrored in the emissions suggests that, in both cases, the primary source of emission for these volatiles are the leaf storage pools, and that the chemotypes of wild cotton plants could differ in their airborne interactions with their environment. Volatile terpenes likely released from stored pools have been found to repel adult moths and piercing-sucking pests of cultivated cotton [86,87], and appear to function as foraging cues for some species of parasitoid wasps [87–90]. Additional research is needed to evaluate how the two chemotypes may impact wild cotton’s interactions with its insect herbivores and their natural enemies. Our results suggest that the difference between the chemotypes may be of lesser importance in this context, as γ-terpinene exhibited a notably low inducibility in comparison to other leaf monoterpenes. Measurements at additional time points and molecular studies would be required to confirm how the synthesis of γ-terpinene and correlated compounds respond to *S. exigua* herbivory. Moreover, as induction of volatiles in cotton can be specific to the species of an attacking herbivore [91–94], we recommend that future work investigates the response of wild cotton chemotypes to other herbivore species.

Increased production of monoterpenes has also been reported in several plant species in response to abiotic stresses such as drought and excessive temperatures, against which they may offer protective properties [7,95–98]. Interestingly, some plant species specifically increase γ-terpinene concentrations in their tissues under drought conditions [99,100], and it has been demonstrated that γ-terpinene has a particularly high antioxidant capacity [101,102]. Previous work on Yucatanese wild cotton documented that populations in higher-temperature environments tend to exhibit a greater density of glands (albeit non-significantly), while herbivory pressure and gland density are not correlated among populations [103]. These findings suggests that abiotic conditions could potentially be significant factors driving variation in gland density and their chemical content. Considering the particularly harsh environments in which wild cotton grows [68], it would be interesting to investigate the relationship between γ-terpinene content and tolerance to extreme environmental conditions.

### 2 Effects of herbivory on the leaf-stored and emitted volatile profiles

Two days of herbivory on the first and second leaves led to a rapid accumulation of several volatile terpenes in the undamaged fourth leaf of the same plants. The leaf terpenes differed in their inducibility, which resulted in compositional changes in leaf volatile profiles triggered by herbivory. A similar variation in inducibility among compounds has previously been described for domesticated varieties, with (*E*)*-*βocimene being the most responsive to damage, and with sesquiterpenes exhibiting a stronger response than monoterpenes [47,48]. The higher inducibility of (*E*)*-*β-ocimene may allow for the increased synthesis of heliocides H1 and H4, the most inducible gland-stored, non-volatile defensive terpenoid aldehydes produced by cotton, for which (*E)-*β-ocimene is a precursor, together with hemigossypolone [47,104,105].

The constitutive volatile emissions of wild cotton consisted mostly of terpenes that also accumulated in the leaves: several cyclic terpenes, along with the linear terpenes β-myrcene and (*E*)*-*β-ocimene. This parallels previous descriptions of cultivated varieties, where these compounds are known to be stored in glands and constitutively released in small amounts [46,49,52]. For most of these terpenes, we found a small increase in their emission rates following herbivory. As there was no active damage occurring on the plant during sampling, this increase could reflect passive release from storage pools in induced undamaged leaves, although we cannot exclude some leakage from very fresh wounds, nor can a release from sources other than storage pools be fully ruled out [9,106]. The latter is already known for (*E*)*-*β-ocimene in cultivated cotton, which is not only accumulated in the leaves but is also *de novo* synthesized and systemically emitted upon induction [52]. Consistent with this previously observed phenomenon, we found a particularly strong induction of (*E*)*-*β-ocimene emissions in comparison to the other terpenes detected in both leaves and headspace. Together with (*E*)*-*β-ocimene, the major compounds induced by *S. exigua* herbivory in wild cotton plants included indole, (*Z*)*-*3-hexenyl acetate, and several terpenes that were distinct from those found in stored pools, namely the linear terpenoids linalool, (*E*)*-*β-farnesene, (*E,E*)*-*α-farnesene, DMNT, and TMTT. This aligns with previous findings in cultivated cotton, showing that these compounds are newly synthesized following damage [49–53,55].

We identified several compounds that, to the best of our knowledge, have not yet been reported for cotton, including aliphatic aldoximes, nitriles, and the aromatic compound 2-phenylethanol. A nitrile compound previously described in wild cotton from Yucatán [58] likely corresponds to the nitrile we could not conclusively identify. Aldoximes are nitrogenous compounds synthesized from amino acids by cytochrome P450 monooxygenases of the CYP79 family, found across all angiosperms, and are precursors to nitriles [107,108]. Aldoximes and nitriles may be precursors to volatile alcohols, such as 2-phenylethanol [108,109]. Various plant species emit volatile aliphatic aldoximes in response to herbivory or jasmonic acid treatment [108]. In *Populus trichocarpa,* herbivore-induced CYP79 enzymes produce (*E/Z*)-2-methylbutyraldoximes from L-isoleucine in response to caterpillar herbivory [107], and CYP71 enzymes convert aldoximes into nitriles, including benzyl nitrile, from the semi-volatile compound phenylacetaldoxime [109]. Volatile aldoximes and nitriles could be involved in cotton indirect defenses; in *Populus nigra*, methylbutyraldoximes and benzyl nitrile are central to the recruitment of parasitoid wasps, despite being only minor components of the full induced volatile blend [110]. Benzyl nitrile is involved in direct defense mechanisms of *Populus trichocarpa and P. nigra* [109,111]. Aldoximes and nitriles might be emitted only in small amounts by cultivated *G. hirsutum*, and therefore more difficult to detect with classic sampling and analyses [112]. Our results could imply that wild cotton plants produce more aldoximes and nitriles than cultivated varieties. This might be of interest for future studies as it could offer an opportunity to enhance the resistance of cultivated cotton varieties to herbivorous pests.

### 3 Variation in the induction of leaf-stored and emitted volatiles

In terms of leaf volatile content, the two chemotypes varied in their inducibility (magnitude of the increase in leaf concentrations) of the monoterpene (*E*)*-*β-ocimene, whereas there was no difference between chemotypes in the induction of (*E*)*-*β-ocimene in the headspace emissions. This is likely due to the ealier mentioned possibility that most of its induced emissions may derive from new synthesis rather than stored pools [52]. This variation in the inducibility of (E)-β-ocimene in leaves could have implications for wild cotton plants’ airborne interactions, particularly in scenarios involving subsequent leaf damage, as it would lead to the release of the induced stored pools [51,53]. As for headspace HIPV emissions, we observed considerable intraspecific variation at the population and regional levels in the emissions of the terpenes (*E,E*)*-*α-farnesene and (*E*)-myroxide, as well as in the emissions of aldoximes. Nitriles were also found to differ, to a lesser extent, among regions. Quantities of aldoximes and nitriles were also observed to vary between the two leaf-terpene chemotypes. As the prevalence of the two chemotypes was partially nested within region, the source of variation in emissions of these compounds cannot be conclusively ascertained. Having said this, the similarity between damaged plants from Chicxulub (high proportion of chemotype B) and Coloradas (dominance of chemotype A) in the production of both classes of compounds suggests that this variation is more likely linked to interpopulation differences than to the chemotypes themselves. In contrast, chemotype-specific responses to caterpillar herbivory have been described in another species (*Tanacetum vulgare*) for both leaf-stored and emitted volatiles [113]. Despite being confined to coastal habitats, Yucatanese wild cotton is subjected to substantial spatial variability in herbivore pressure and abiotic conditions [103,114–116], which could be associated with the origin of the observed variation in HIPVs. Further work is required to understand the drivers of these differences. Given the multiple proposed functions of HIPVs in plant defense against insect herbivores [2], the differences in HIPVs that we observed could have consequences for wild cotton’s resistance to herbivores. Genotypes producing higher amounts of certain HIPVs that enhance their direct or indirect defenses as well as those of neighboring plants [57] could be a valuable genetic resource in the development of more resistant cotton cultivars. The leaf volatiles may also enhance tolerance to abiotic stresses such as extreme temperatures and drought [7], which is worthy of further exploration.

## Conclusions

In this study, we present a comprehensive description of the quantitative and qualitative changes triggered by caterpillar herbivory in the leaf-stored volatile terpenes and compounds emitted by wild *G. hirsutum* plants from the Yucatán Peninsula. We show that two previously described monoterpene chemotypes can also be distinguished based on their headspace emissions, which is true for both undamaged and caterpillar-damaged plants. Furthermore, we provide a first report of aldoximes being identified as cotton HIPVs. Wild cotton volatile responses to herbivory were found to vary among chemotype, geographic location and population. Future research should examine the consequences of this variation for the plant’s resistance to herbivory, or other biotic and abiotic stresses.

## Ethics approval and consent to participate

Not applicable

## Consent for publication

Not applicable

## Availability of data and materials

The datasets generated during the current study, as well as the R code for the analyses are available in the Zenodo repository, https://doi.org/10.5281/zenodo.10927508. The procedure for spectra analyses of GC-MS data is available at https://github.com/bustossc/PyMS-routine.

## Authors’ contributions

T.C.J.T. and L.A-R., C.B-S., M.V.C. and M.M. designed the research. L.A-R., T.Q-M. and B.P-N. collected the cotton seeds. C.B-S., M.V.C., M.M. and G.F. performed the experiment and processed the samples. M.M. and C.B-S. processed and analyzed the data. M.M. wrote the first draft of the manuscript with contributions by all authors.

## Acknowledgments

The authors thank Thomas Degen (UniNE) for the illustrations of cotton and for helping to create figure 1, Karla Ku Durán (UADY) for laboratory support, Dominic Davis-Foster (Staffordshire U) for maintaining the PyMassSpec repository, Tobias Köllner (Max Planck Institute for Chemical Ecology) for the aldoximes standards, and Fanny Deiss, Sara Leite Dias, Jonathan Interian, and Miguel Cárdenas for help with sample collection.

