## Supplemental Material for "Induction by caterpillars of stored and emitted volatiles in terpene chemotypes from populations of wild cotton (*Gossypium hirsutum*)"

#### Supplementary figure 1

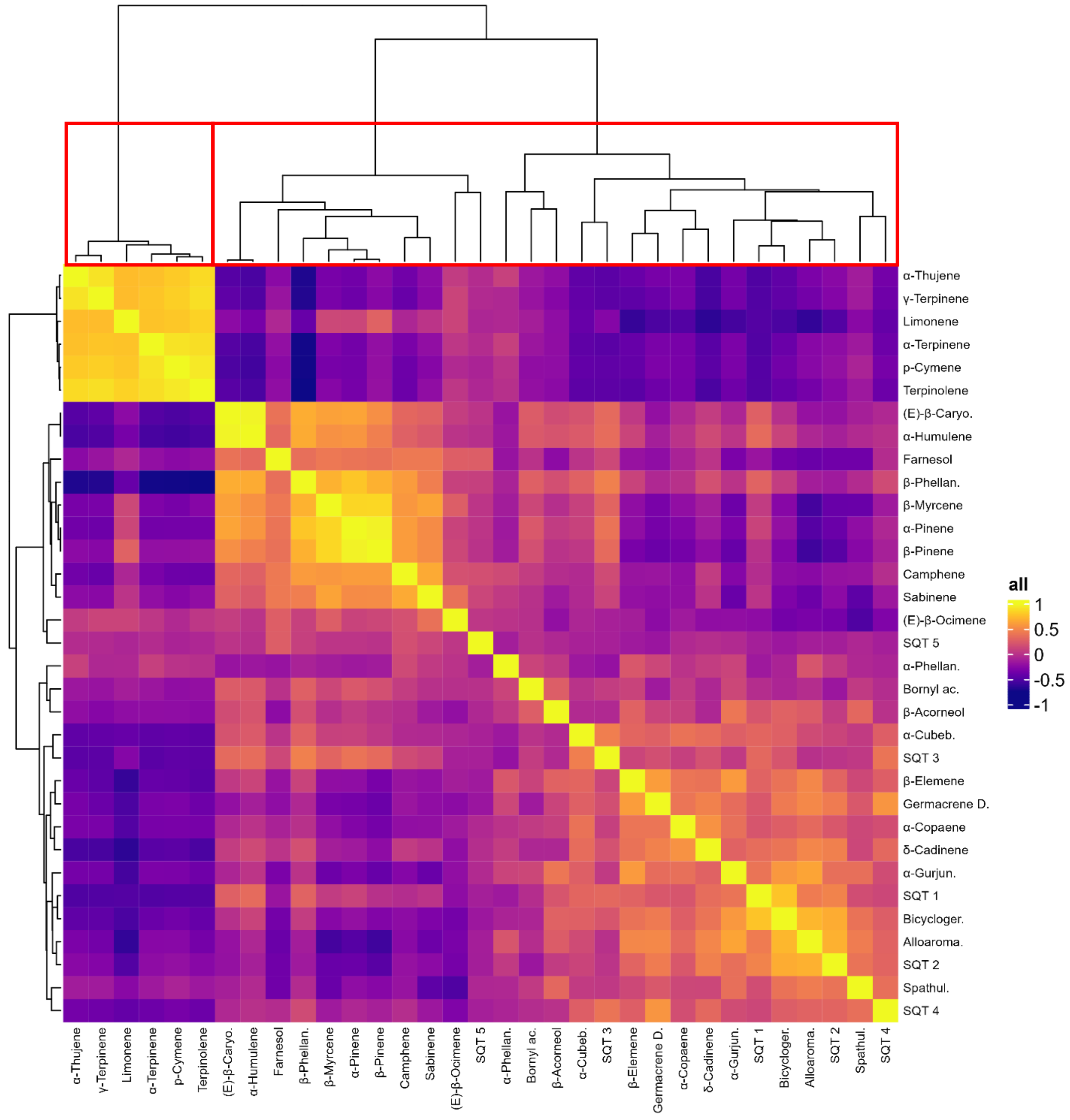


Correlations between relative abundances of stored leaf mono- and sesquiterpenes in undamaged wild cotton plants. Red squares indicate the largest statistically significant clusters (Approximately Unbiased p-values < 0.05).

#### Supplementary figure 2

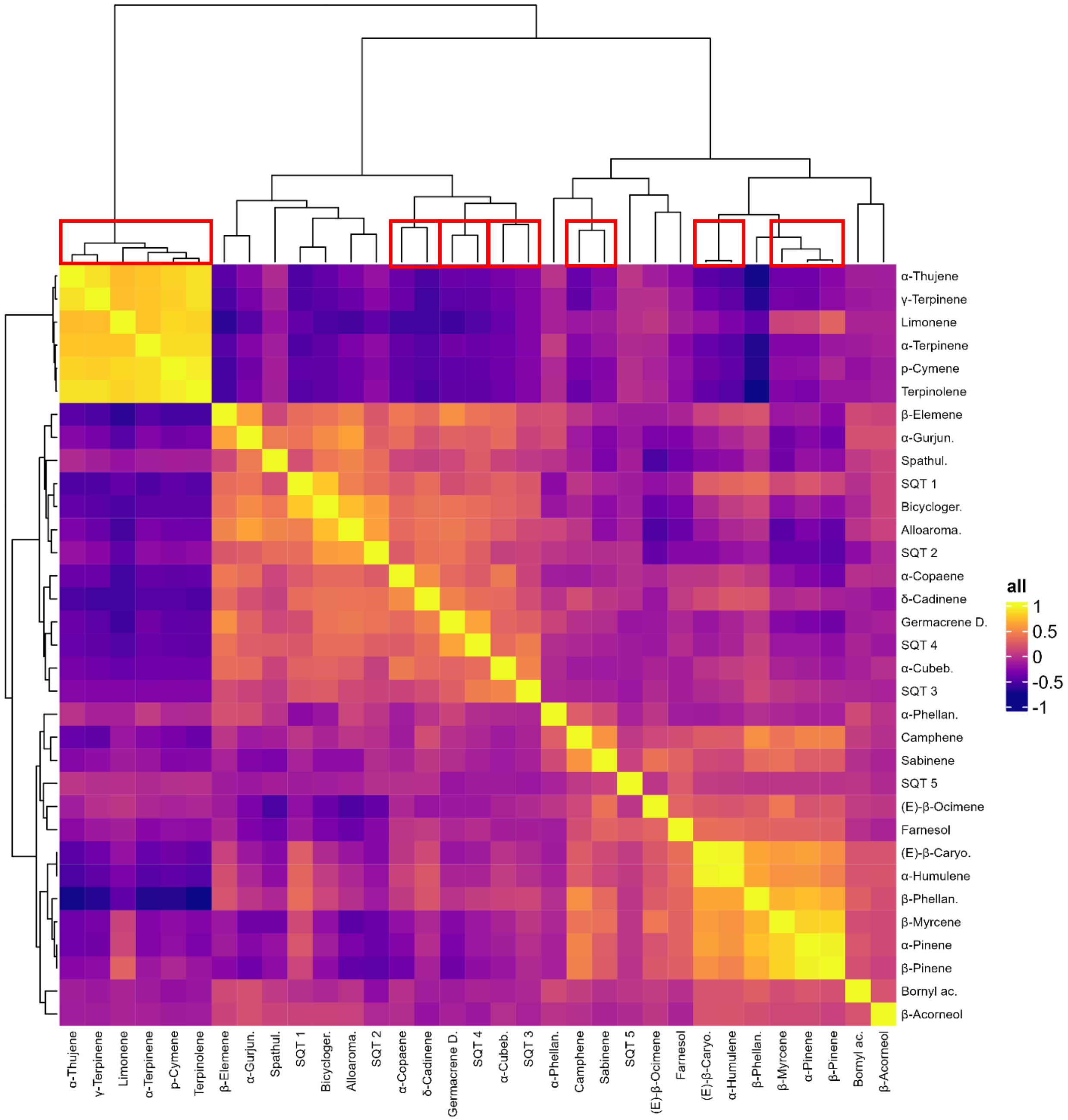


Correlations between relative abundances of stored leaf mono- and sesquiterpenes in wild cotton plants damaged by *Spodoptera exigua*. Red squares indicate the largest statistically significant clusters (Approximately Unbiased p-values < 0.05)

### Supplementary figure 3

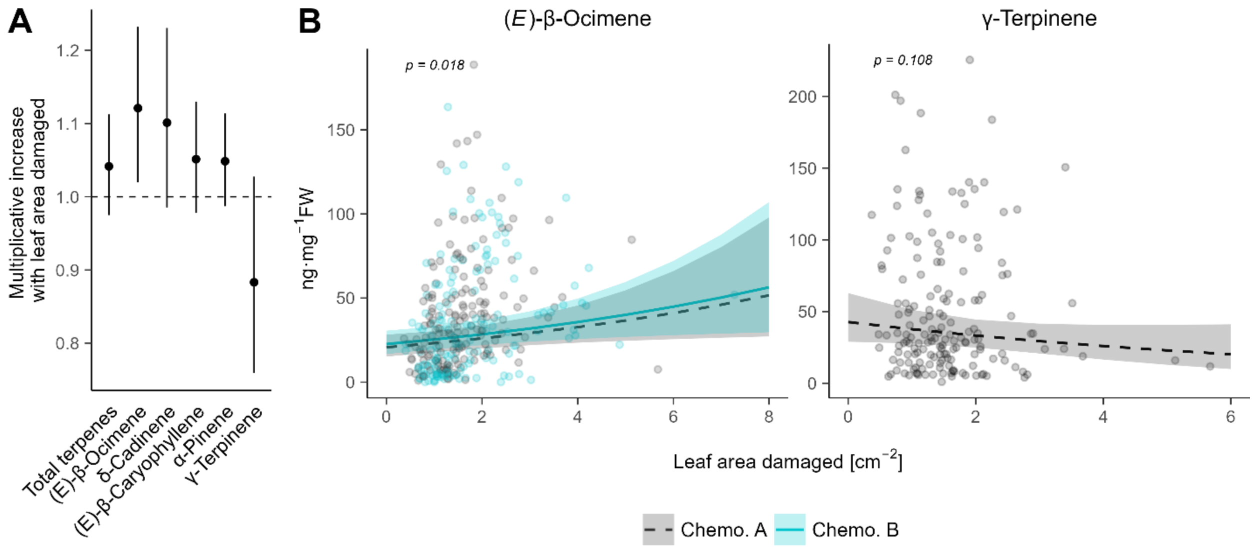


Relationship between damage intensity and accumulation of volatile mono- and sesquiterpenes in leaves of wild cotton. A: multiplicative increases (back-transformed coefficient estimates from GLMMs and their 95% CIs. The dotted line indicates a coefficient of 1, where quantities emitted remain constant with damaged leaf area. B: adjusted predictions and 95% CI from GLMMs with observed values as data points. γ-Terpinene was analysed within chemotype A while all other compounds were analysed for both chemotypes together, with chemotype set as fixed factor.

#### Supplementary figure 4

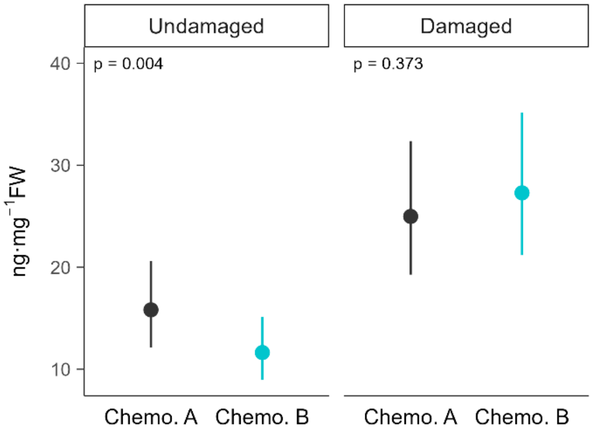


Variation between chemotypes in the leaf concentrations of (*E*)-β-ocimene in undamaged wild cotton and plants damaged by *Spodoptera exigua* for two days. Estimated marginal means and 95% CI from GLMMs performed separately for undamaged and damaged plants. The model on damaged plants includes damage intensity as covariate.

#### Supplementary figure 5

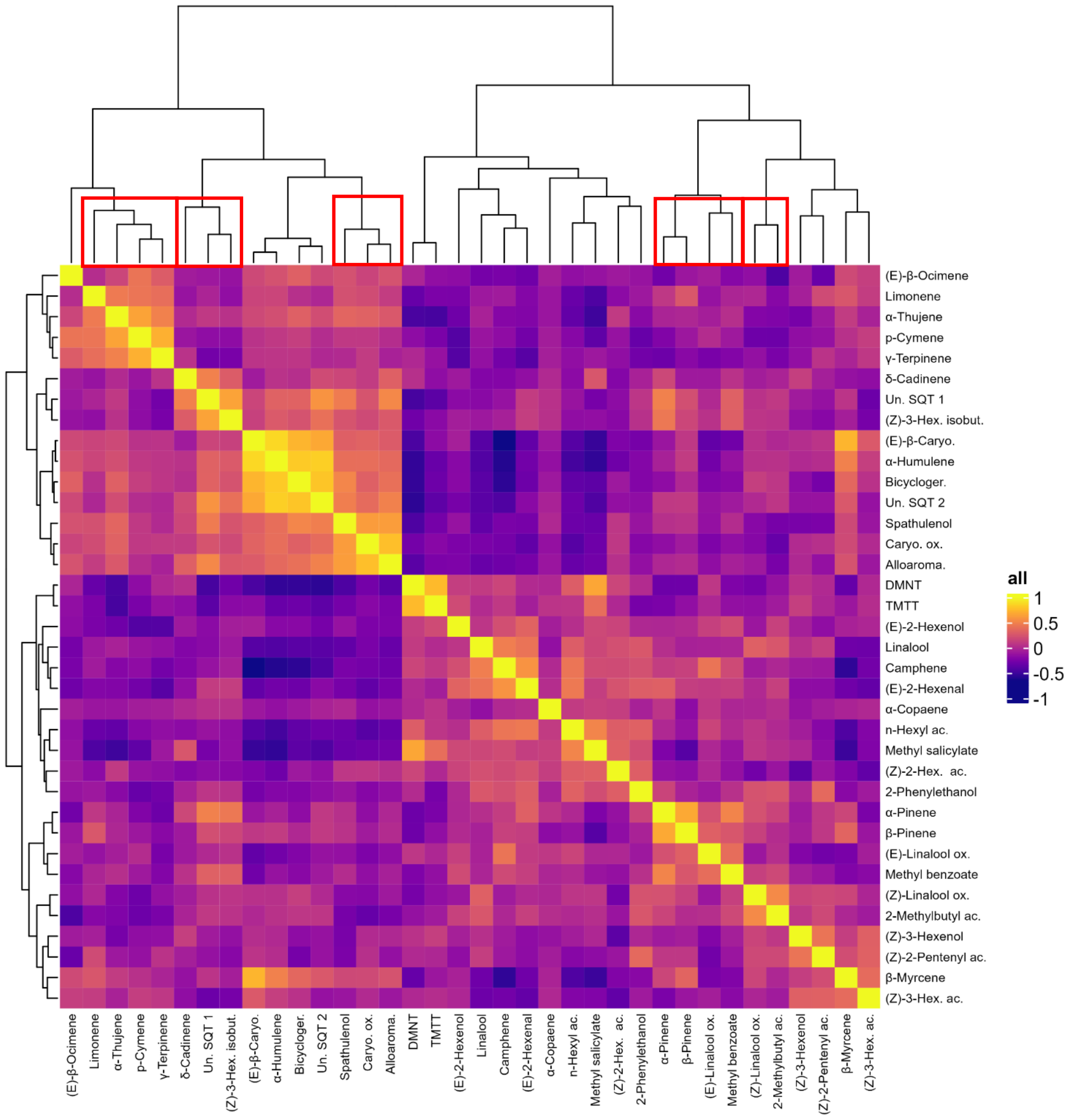


Correlations between relative abundances of all volatiles emitted by undamaged wild cotton plants. Red squares indicate largest statistically significant clusters (Approximately Unbiased p-values < 0.05). Only compounds at least in 10% of the undamaged plants were considered.

#### Supplementary figure 6

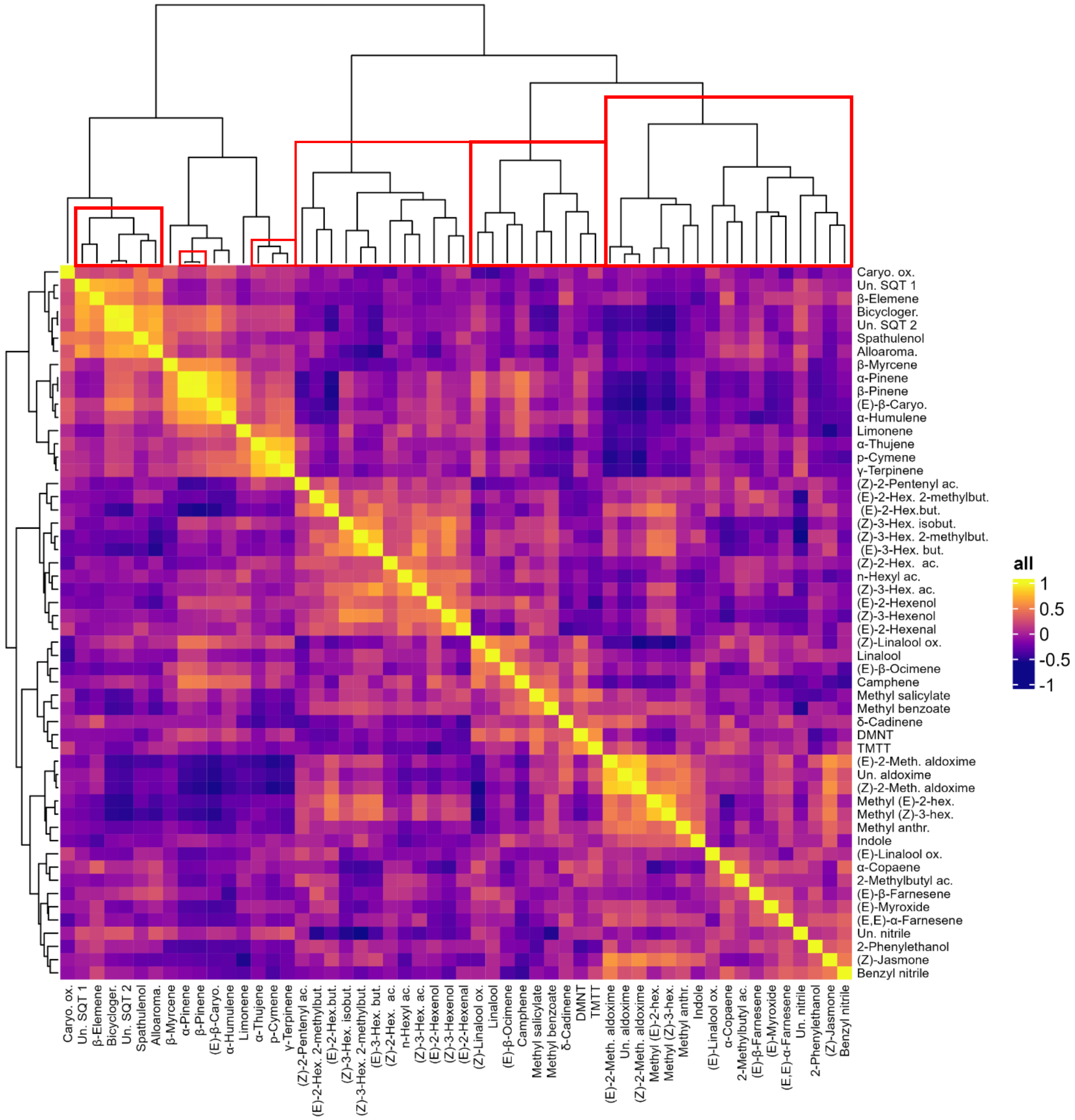


Correlations between relative abundances of all volatiles emitted by wild cotton plants damaged by *Spodoptera exigua*. Red squares indicate largest statistically significant clusters (Approximately Unbiased p-values < 0.05).

### Supplementary figure 7

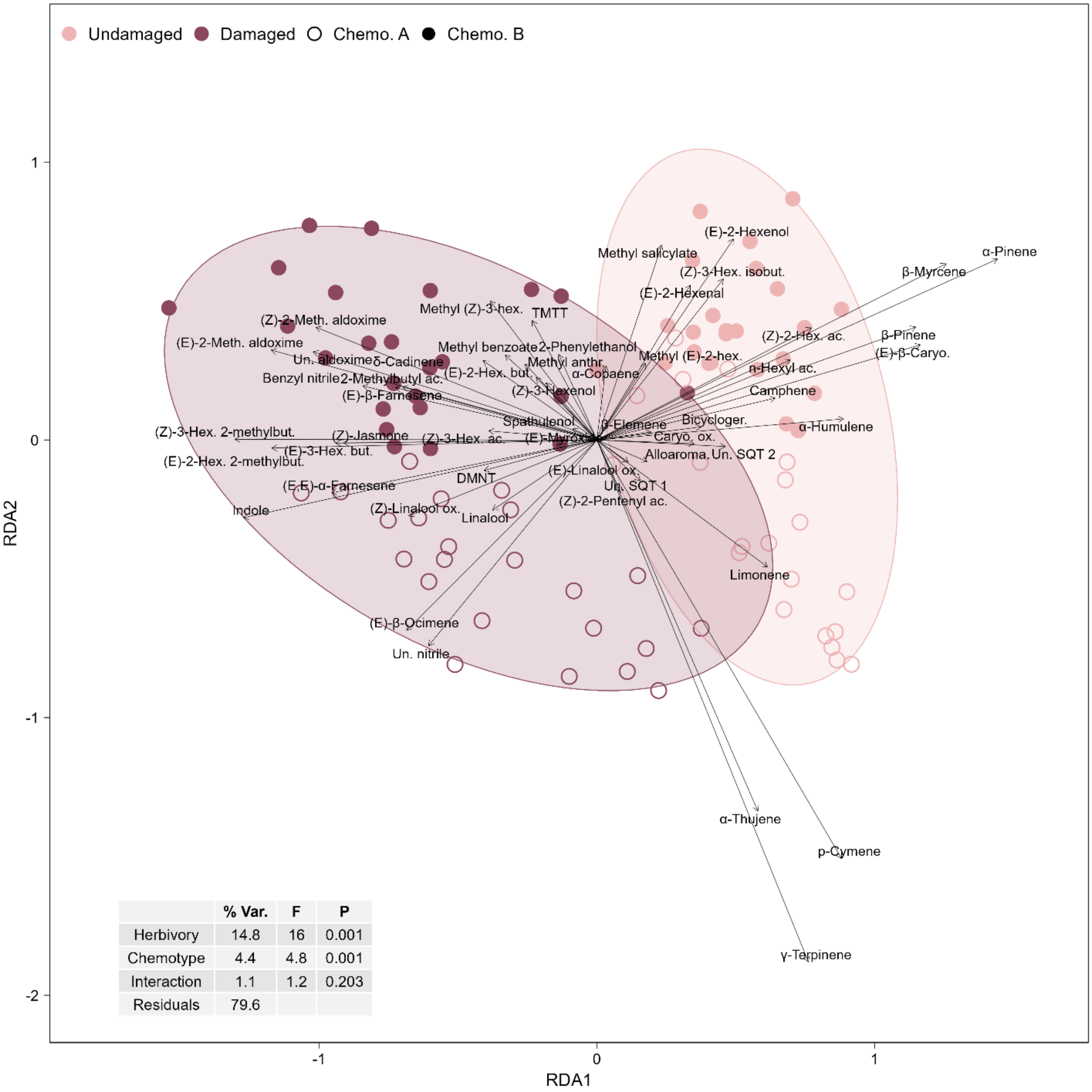


Biplot with symmetric scaling of a pRDA on the relative abundances (clr-transformed) of all compounds quantified in the headspace samples. Highly inducible compounds were defined as the ones with a negative score on the first axis.

#### Supplementary figure 8

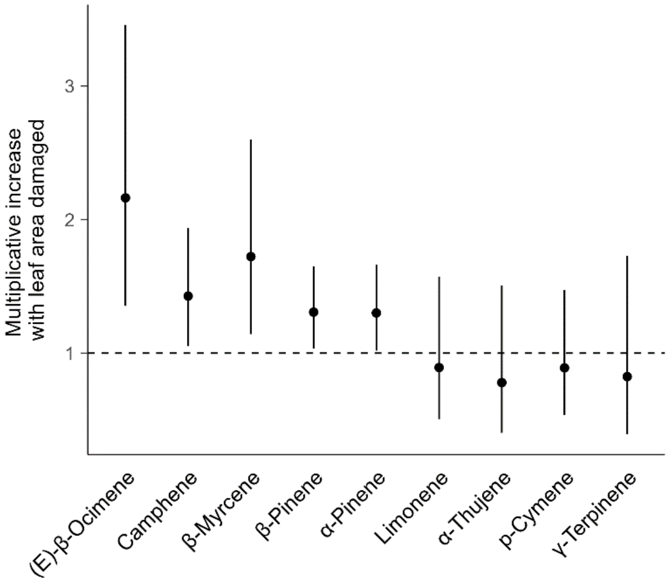


Relationship between damage intensity and emission rates of monoterpenes detected in both headspace and leaf extracts. Multiplicative increases with the leaf area damaged (back-transformed coefficient estimates of GLMMs and their 95% CIs). The dotted line indicates a coefficient of 1, where quantities emitted remain constant with damaged area. γ-Terpinene, p-cymene, α-thujene and limonene were analyzed within chemotype A while all other compounds were analyzed for both chemotypes together, with chemotype set as fixed factor.

#### Supplementary figure 9

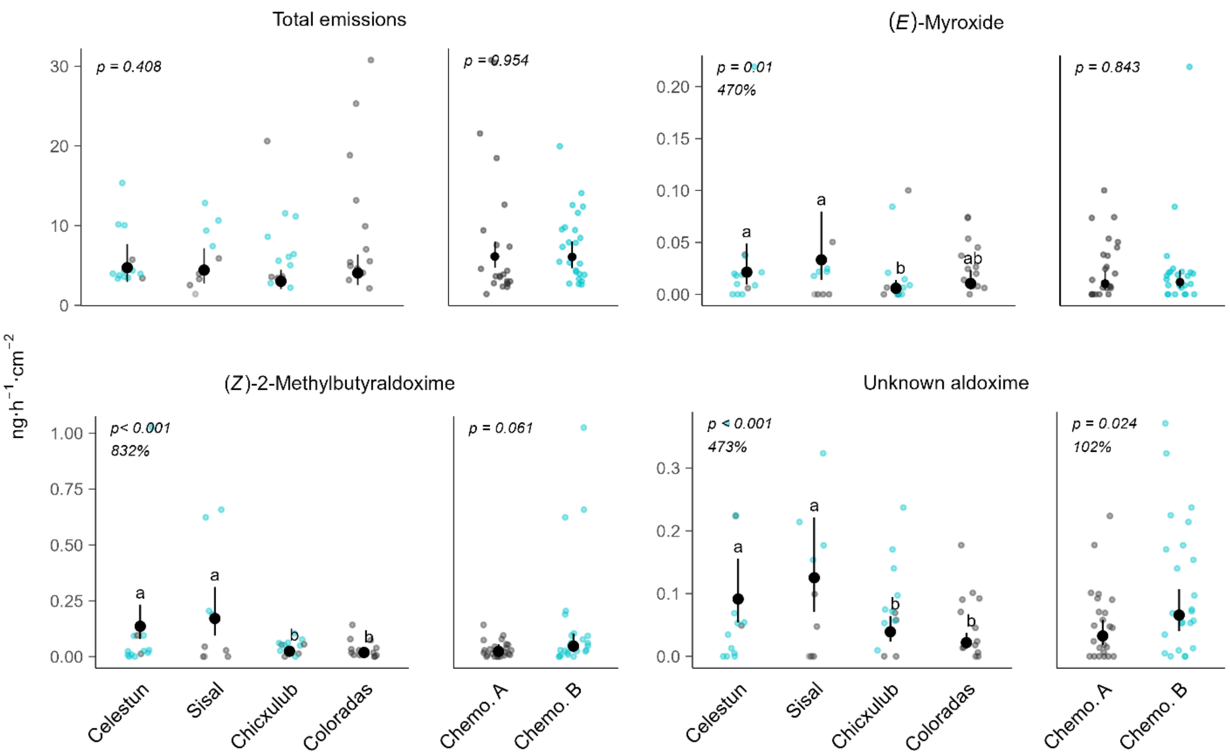


Variation in total emissions of herbivore-induced volatiles (highly induced compounds only) and in emissions of (*E*)-myroxide and aldoximes among the four regions grouping wild cotton populations along the Yucatan coast and between the two leaf-terpene chemotypes. Headspace volatile emissions were sampled from 48 wild cotton plant damaged by *Spodoptera exigua* for two days. Regions are displayed from west to east. Prevalence of chemotypes is nested with regions, as indicated by the color of data points (grey points indicate chemotype A; blue points indicate chemotype B), therefore they were analyzed separately. Estimated marginal means and 95% CI from GLMMs with observed values as data points. Damage intensity was controlled for as covariate. Different letters indicate significant pairwise comparisons after multiple test adjustment (FDR). For regions, % values indicate the difference between the region with the lowest and highest estimated mean emission rates.

### Supplementary figure 10

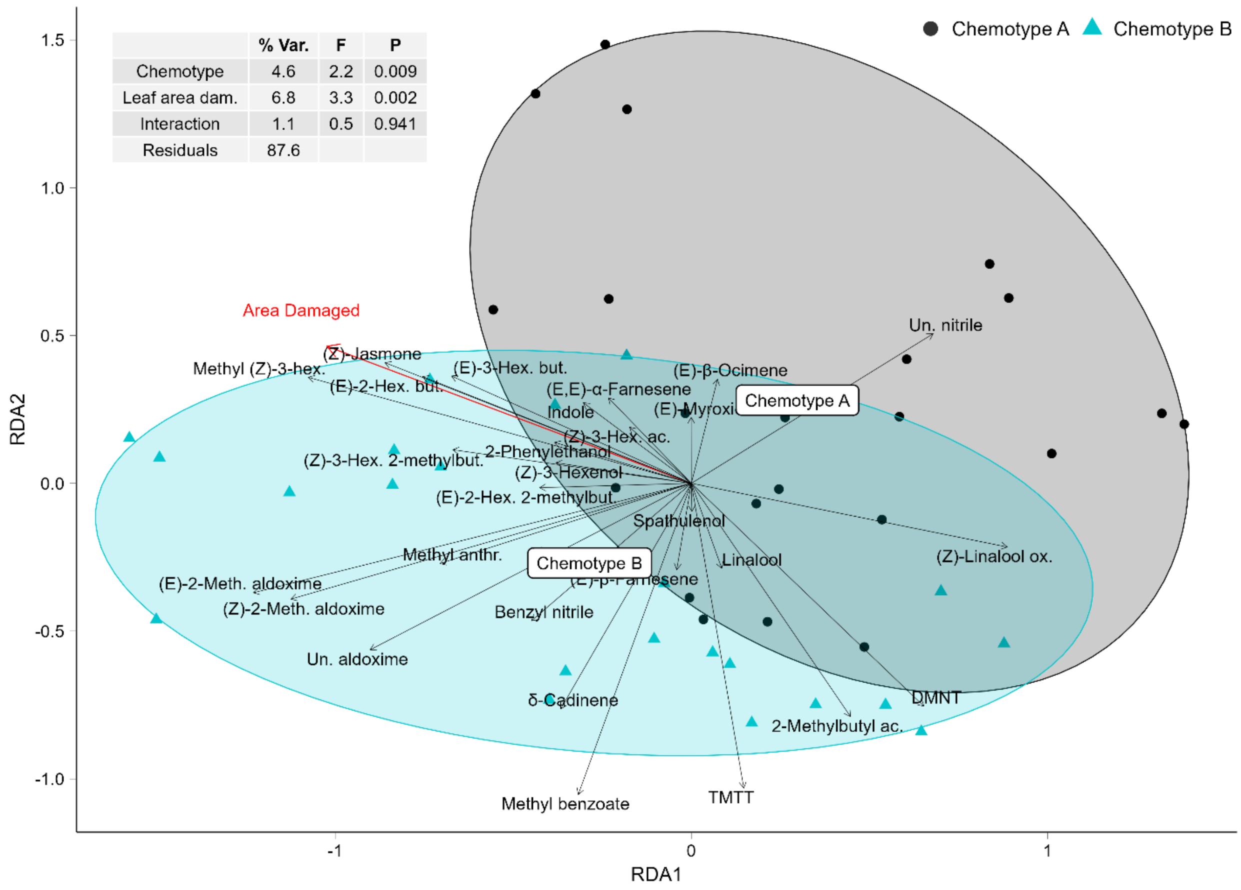


| Differences in emission rates of damaged plants between the two chemotypes | |
|---|---|
| | **Chemotype** |
| **Unknown nitrile** | $\chi^2_{(1)} = 7.100$, **p = 0.008** |
| **TMTT** | $\chi^2_{(1)} = 5.971$, **p = 0.014** |
| Methyl benzoate | $\chi^2_{(1)} = 2.970$, p = 0.085 |
| δ-Cadinene | $\chi^2_{(1)} = 0.450$, p = 0.502 |
| **Benzyl nitrile** | $\chi^2_{(1)} = 4.709$, **p = 0.039** |
| Methyl anthranilate | $\chi^2_{(1)} = 1.129$, p = 0.288 |
| **Unknown aldoxime** | $\chi^2_{(1)} = 5.069$, **p = 0.024** |
| **(*Z*)-2-methylbutyraldoxime** | $\chi^2_{(1)} = 3.503$, p = 0.061 |
| **(*E*)-2-methylbutyraldoxime** | $\chi^2_{(1)} = 7.237$, **p = 0.007** |
| **Inducible compounds differing between chemotypes but detected in undamaged plants: variation in the change with herbivory** | |
| | **Chemotype x Herbivory** |
| TMTT | $\chi^2_{(1)} = 1.335$, p = 0.248 |

Biplot with symmetric scaling of a pRDA on the abundances of highly inducible compounds relative to the whole volatile blend emitted by wild cotton plants damaged by *Spodoptera exigua* (clr-transformed). Highly inducible compounds were defined as described in supplementary figure 7. All compounds are plotted. The table is a summary of the GLMMs analyzing the effect of chemotype on the emission rates of compounds highlighted as potentially varying between chemotypes by the pRDA. Damage intensity was controlled for as covariate in models performed on damaged plants. Chi-squared ($\chi^2$) values are shown, and statistically significant ($p < 0.05$) effects are indicated in bold.

#### Supplementary figure 11

Biplot with symmetric scaling of a pRDA on the abundances of highly inducible compounds relative to the whole volatile blend emitted by wild cotton plants damaged by *Spodoptera exigua* (clr-transformed). Highly inducible compounds were defined as described in supplementary figure 7. The table is a summary of the GLMMs analyzing the effect of region on the emission rates of compounds highlighted as potentially varying between chemotypes by the pRDA. Damage intensity was controlled for as covariate in models performed on damaged plants. Chi-squared ($\chi^2$) values are shown, and statistically significant ($p < 0.05$) effects are indicated in bold.

| **Differences in emission rates of damaged plants among the four regions** | |
|---|---|
| | **Region** |
| **Unknown nitrile** | $\chi^2_{(3)} = 8.034$, **p = 0.045** |
| (*Z*)-Linalool oxide | $\chi^2_{(3)} = 2.400$, p = 0.493 |
| DMNT | $\chi^2_{(3)} = 0.916$, p = 0.821 |
| TMTT | $\chi^2_{(3)} = 3.386$, p = 0.335 |
| (*E*)-β-Ocimene | $\chi^2_{(3)} = 1.284$, p = 0.733 |
| Linalool | $\chi^2_{(3)} = 3.632$, p = 0.304 |
| **δ-Cadinene** | $\chi^2_{(3)} = 14.293$, **p = 0.002** |
| Methyl benzoate | $\chi^2_{(3)} = 7.670$, p = 0.053 |
| **Unknown aldoxime** | $\chi^2_{(3)} = 25.789$, **p < 0.001** |
| **(*Z*)-2-methylbutyraldoxime** | $\chi^2_{(3)} = 71.290$, **p < 0.001** |
| **(*E*)-2-methylbutyraldoxime** | $\chi^2_{(3)} = 62.022$, **p < 0.001** |
| (*E*)-3-Hexenyl butyrate | $\chi^2_{(3)} = 3.880$, p = 0.274 |
| (Z)-3-Hexenyl 2-methylbutyrate | $\chi^2_{(3)} = 3.928$, p = 0.269 |
| **(*E,E*)-α-Farnesene** | $\chi^2_{(3)} = 25.448$, **p < 0.001** |
| (*E*)-β-Farnesene | $\chi^2_{(3)} = 1.823$, p = 0.610 |
| **(*E*)-β-Myroxide** | $\chi^2_{(3)} = 11.442$, **p = 0.009** |
| Spathulenol | $\chi^2_{(3)} = 6.679$, p = 0.083 |
| **Inducible compounds differing among regions but detected in undamaged plants: variation in the increase with herbivory** | |
| | **Region x Herbivory** |
| δ-Cadinene | $\chi^2_{(3)} = 4.809$, p = 0.186 |

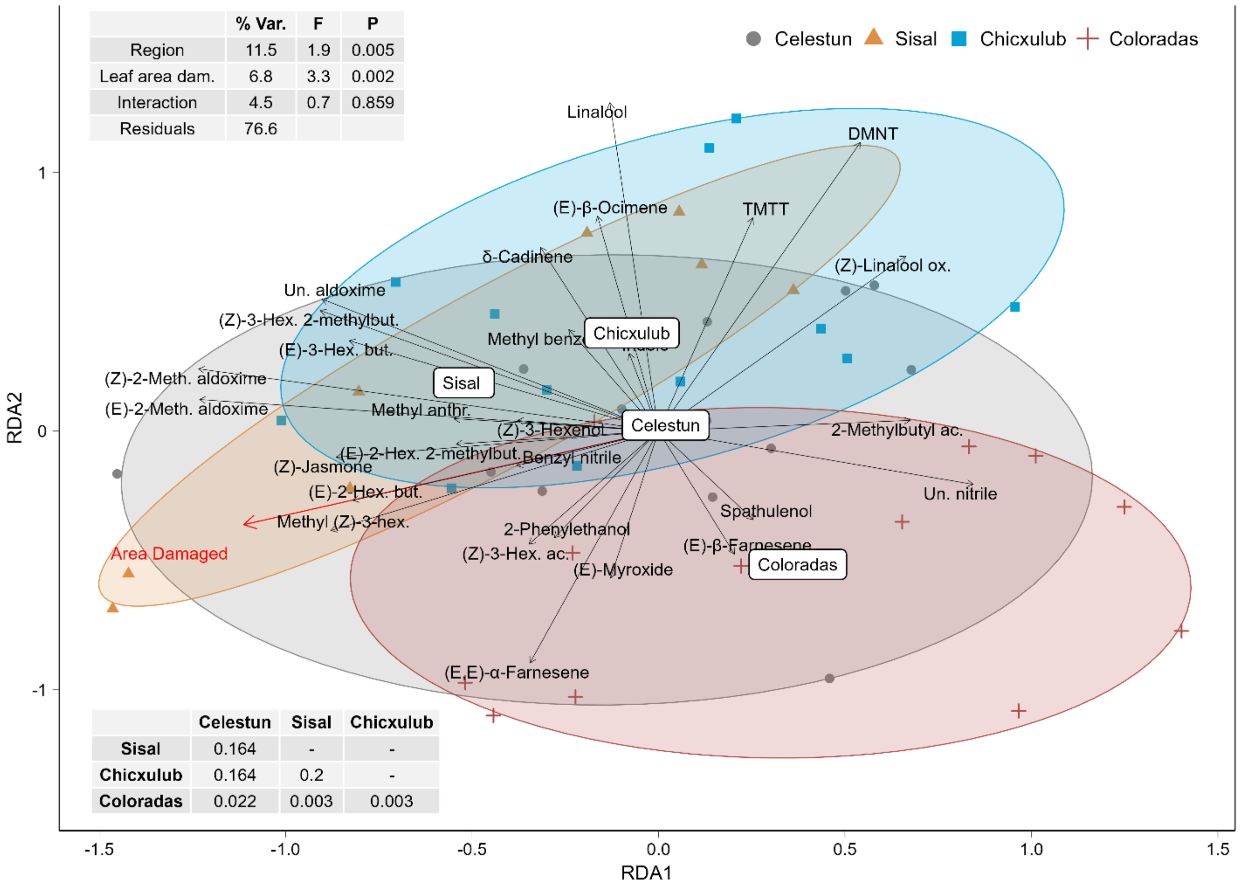

#### Supplementary Table 1

Information about the sites of seeds collection and number of replicates

| | Leaf extracts | | Headspace | |
|---|---|---|---|---|
| **Region** | **Undamaged # Plants** | **Damaged # Plants** | **Undamaged # Plants** | **Damaged # Plants** |
| Celestún | 86 | 79 | 11 | 13 |
| Sisal | 71 | 60 | 9 | 9 |
| Chicxulub | 80 | 70 | 12 | 13 |
| Coloradas | 116 | 113 | 13 | 13 |
| **Sum** | 353 | 322 | 45 | 48 |

| | Leaf extracts | | Headspace | | | | Leaf extracts | Headspace |
|---|---|---|---|---|---|---|---|---|
| **Population** | **Undamaged #Plants** | **Damaged #Plants** | **Undamaged #Plants** | **Damaged #Plants** | **Latitude** | **Longitude** | **# Genotypes/Population** | **# Genotypes/Population** |
| Drilos (DR) | 20 | 19 | 1 | 4 | 20.94668 | -90.36220 | 5 | 3 |
| Vergel (VR) | 21 | 20 | 3 | 3 | 20.97018 | -90.35161 | 5 | 3 |
| Consorcio (CN) | 21 | 20 | 3 | 2 | 20.99828 | -90.33714 | 6 | 3 |
| Chapo (CH) | 24 | 20 | 4 | 4 | 21.01394 | -90.32414 | 6 | 3 |
| Palapita (PL) | 15 | 9 | 4 | 4 | 21.17525 | -89.99657 | 4 | 2 |
| Preseco (PS) | 21 | 19 | 2 | 2 | 21.19097 | -89.96278 | 6 | 2 |
| Seco (SC) | 28 | 26 | 0 | 1 | 21.19306 | -89.95950 | 6 | 1 |
| Papadzul (PP) | 7 | 6 | 3 | 2 | 21.20333 | -89.94119 | 3 | 3 |
| Dos Torres (DT) | 16 | 12 | 3 | 3 | 21.29613 | -89.57930 | 6 | 3 |
| Pre-Tropical (PT) | 22 | 21 | 2 | 3 | 21.30261 | -89.54806 | 6 | 3 |
| Tropical Riviera (TR) | 21 | 18 | 3 | 3 | 21.30396 | -89.54164 | 6 | 3 |
| Cinvestav (CV) | 21 | 19 | 4 | 4 | 21.34116 | -89.30483 | 5 | 4 |
| Holchit (HC) | 29 | 28 | 2 | 2 | 21.60842 | -88.05024 | 6 | 2 |
| Sin Perros (SP) | 30 | 29 | 3 | 3 | 21.60858 | -87.96481 | 6 | 3 |
| Cangrejos (CJ) | 29 | 30 | 5 | 5 | 21.59358 | -87.91953 | 6 | 3 |
| Chorlitos (CO) | 28 | 26 | 3 | 3 | 21.58556 | -87.89372 | 6 | 3 |

#### Supplementary table 2

Structures of the GLMMs used to analyze concentrations of volatile terpenes in wild cotton leaves. Chemo = chemotype, Area = fourth leaf area, Damage = damage intensity (total leaf area consumed by *S. exigua*). Full fixed effects are shown, interaction terms were removed if not significantly contributing to the model.

| Data | Model | Effects tested with the model | Fixed effects | Random effects (intercepts and slopes) |
|---|---|---|---|---|
| All plants | A | Herbivory / Chemotype / Chemotype-by-Herbivory interaction | Herbivory + Chemo + Chemo:Herbivory + Area + Batch + Area:Batch | Population + Genotype |
| | B | Genotype-by-Herbivory interaction | Herbivory + Chemo + Chemo:Herbivory + Area + Batch + Area:Batch | Population + Herbivory:Genotype |
| | C | Population-by-Herbivory interaction | Herbivory + Chemo + Chemo:Herbivory + Area + Batch + Area:Batch | Herbivory: Population + Genotype |
| | D | Region-by-Herbivory interaction | Herbivory + Region + Region:Herbivory + Area + Batch + Area:Batch | Population + Genotype |
| Damaged plants | E | Damage intensity | Damage + Chemo + Chemo:Damage + Area + Batch + Area: Batch | Population + Genotype |

#### Supplementary table 3

Structures of the GLMMs used to analyze emission rates of volatiles by wild cotton plants. Chemo = chemotype, Area = fourth leaf area, Damage = damage intensity (total leaf area consumed by *Spodoptera exigua*). Full models are shown, interaction terms were removed if not significantly contributing to the model.

| Data | Model | Effects tested with the model | Structure of the model | |
|---|---|---|---|---|
| | | | **Fixed effects** | **Random effects (intercepts)** |
| **All plants** | A | Herbivory / Chemotype-by-Herbivory interaction | Herbivory + Chemo + Chemo:Herbivory + Area + Batch + Area:Batch | Population + Genotype |
| | B | Region-by-Herbivory interaction | Herbivory + Region + Region:Herbivory + Area + Batch + Area:Batch | Population + Genotype |
| **Damaged plants** | C | Chemotype / Damage intensity | Chemo + Damage + Chemo:Damage + Area + Batch + Area:Batch | Population |
| | D | Region | Region + Damage + Region:Damage + Area + Batch + Area:Batch | Population |
| | E | Population | Chemo + Damage + Chemo:Damage + Area + Batch + Area:Batch | Population |

#### Supplementary table 4

Volatiles quantified in the leaves of undamaged and damaged wild cotton plants (>10% of samples). Median leaf concentrations and quantiles. RI: retention index. RRF: standards used for quantification by relative response factor.  Compounds marked with an asterisk were authenticated via comparison to standards.

| | | | **Undamaged plants ng mg$^{-1}$ FW** | | | | | **Damaged plants ng mg$^{-1}$ FW** | | | | |
|---|---|---|---|---|---|---|---|---|---|---|---|---|
| **Compounds** | **RI** | **RRF** | **Q10** | **Q25** | **Median** | **Q75** | **Q90** | **Q10** | **Q25** | **Median** | **Q75** | **Q90** |
| **(*Z*)-3-Hexenal** | 797 | (*Z*)-3-Hexenol | 33.189 | 48.239 | 64.875 | 81.373 | 98.537 | 71.119 | 87.934 | 104.141 | 123.76 | 141.217 |
| **(*E*)-2-Hexenal** | 849 | (*Z*)-3-Hexenol | 2.436 | 3.935 | 5.524 | 7.525 | 8.776 | 5.407 | 6.733 | 8.781 | 11.033 | 12.63 |
| **(*E,E*)-2,4-Hexadienal** | 910 | (*Z*)-3-Hexenol | 12.913 | 22.379 | 35.004 | 69.297 | 96.763 | 23.89 | 35.817 | 49.248 | 79.42 | 103.109 |
| **α-Thujene** | 929 | α-Pinene | 0 | 0.488 | 0.999 | 2.241 | 4.517 | 0 | 0 | 1.087 | 2.334 | 6.101 |
| **α-Pinene*** | 938 | α-Pinene | 53.242 | 76.088 | 113.834 | 172.234 | 225.443 | 57.269 | 81.222 | 116.888 | 178.769 | 236.72 |
| **Camphene** | 955 | α-Pinene | 0 | 1.591 | 2.305 | 3.324 | 4.214 | 0 | 1.649 | 2.508 | 3.607 | 4.795 |
| **Sabinene** | 977 | α-Pinene | 2.769 | 4.922 | 6.892 | 9.485 | 13.211 | 0.53 | 5.225 | 7.632 | 10.132 | 14.205 |
| **β-Pinene** | 983 | α-Pinene | 10.521 | 14.325 | 21.083 | 32.488 | 42.822 | 11.443 | 16.046 | 22.252 | 34.582 | 47.464 |
| **β-Myrcene*** | 989 | β-Myrcene | 43.381 | 59.826 | 88.811 | 133.979 | 180.069 | 45.949 | 62.527 | 92.878 | 140.468 | 180.928 |
| **α-Phellandrene** | 1008 | α-Pinene | 0 | 0 | 0.924 | 1.662 | 2.633 | 0 | 0 | 1.155 | 2.188 | 2.921 |
| **α-Terpinene** | 1020 | γ-Terpinene | 0 | 0 | 0 | 1.459 | 3.044 | 0 | 0 | 0 | 1.755 | 3.924 |
| **p-Cymene** | 1028 | γ-Terpinene | 0 | 0 | 0 | 1.385 | 3.216 | 0 | 0 | 0 | 1.787 | 4.288 |
| **Limonene*** | 1033 | Limonene | 3.607 | 5.357 | 8.576 | 16.524 | 30.263 | 4.248 | 6.144 | 9.929 | 17.005 | 39.638 |
| **β-Phellandrene** | 1035 | γ-Terpinene | 0 | 2.075 | 4.175 | 6.455 | 10.341 | 0 | 1.549 | 4.747 | 7.168 | 11.331 |
| **(*E*)-β-Ocimene** | 1047 | γ-Terpinene | 2.62 | 5.381 | 13.118 | 26.512 | 47.123 | 4.259 | 12.729 | 29.694 | 51.903 | 84.456 |
| **γ-Terpinene** | 1063 | γ-Terpinene | 0 | 0 | 3.946 | 32.065 | 63.088 | 0 | 0 | 5.754 | 33.838 | 91.493 |
| **Terpinolene** | 1090 | γ-Terpinene | 0 | 0 | 0 | 2.023 | 4.06 | 0 | 0 | 0 | 2.168 | 5.808 |
| **Bornyl acetate** | 1288 | γ-Terpinene | 0 | 0 | 0 | 0 | 2.030 | 0 | 0 | 0 | 0 | 2.511 |
| **Unknown SQT 1** | 1347 | (E)-β-Caryophyllene | 1.566 | 2.542 | 4.899 | 10.803 | 15.636 | 1.892 | 3.225 | 5.869 | 11.153 | 17.759 |
| **α-Cubebene** | 1359 | (*E*)-β-Caryophyllene | 0 | 0 | 0 | 0 | 1.246 | 0 | 0 | 0 | 0 | 2.02 |
| **α-Copaene** | 1389 | (*E*)-β-Caryophyllene | 0 | 0 | 1.477 | 3.498 | 5.383 | 0 | 0 | 2.253 | 4.648 | 8.343 |
| **β-Elemene** | 1401 | (*E*)-β-Caryophyllene | 0 | 0 | 1.245 | 2.563 | 3.787 | 0 | 0 | 1.577 | 3.266 | 4.511 |
| **α-Gurjunene** | 1425 | (*E*)-β-Caryophyllene | 0 | 0 | 0.748 | 2.19 | 3.746 | 0 | 0 | 0.82 | 2.637 | 4.864 |
| **(*E*)-β-Caryophyllene*** | 1432 | (*E*)-β-Caryophyllene | 51.31 | 90.028 | 166.794 | 291.225 | 387.488 | 66.058 | 112.735 | 186.548 | 321.953 | 437.806 |
| **Alloaromadendrene** | 1453 | (*E*)-β-Caryophyllene | 0 | 0 | 2.876 | 7.954 | 12.19 | 0 | 0 | 3.598 | 8.396 | 12.843 |
| **α-Humulene** | 1465 | (*E*)-β-Caryophyllene | 13.338 | 22.615 | 44.039 | 79.025 | 108.515 | 16.334 | 28.611 | 49.841 | 90.095 | 125.651 |
| **Unknown SQT 2** | 1473 | (*E*)-β-Caryophyllene | 0 | 0.633 | 1.472 | 4.444 | 7.977 | 0 | 0.908 | 1.753 | 4.28 | 8.045 |
| **Bicyclogermacrene** | 1498 | (*E*)-β-Caryophyllene | 8.882 | 17.63 | 39.568 | 108.677 | 165.616 | 11.89 | 23.769 | 49.621 | 110.859 | 185.77 |
| **β-Acorneol** | 1519 | (*E*)-β-Caryophyllene | 0 | 0 | 0 | 0.464 | 0.974 | 0 | 0 | 0 | 0 | 0.807 |
| **δ-Cadinene** | 1524 | (*E*)-β-Caryophyllene | 0 | 1.136 | 2.401 | 4.196 | 6.711 | 0 | 1.663 | 3.051 | 5.596 | 8.131 |
| **Germacrene D-4-ol** | 1603 | (*E*)-β-Caryophyllene | 0 | 0 | 0 | 2.209 | 3.512 | 0 | 0 | 1.171 | 2.978 | 4.743 |
| **Spathulenol** | 1610 | (*E*)-β-Caryophyllene | 0 | 0 | 0 | 1.412 | 3.644 | 0 | 0 | 0 | 1.671 | 3.259 |
| **Unknown SQT 3** | 1628 | (*E*)-β-Caryophyllene | 0 | 0 | 0 | 0 | 2.135 | 0 | 0 | 0 | 0 | 2.37 |
| **Unknown SQT 4** | 1668 | (*E*)-β-Caryophyllene | 0 | 0 | 0 | 2.339 | 4.412 | 0 | 0 | 0 | 2.971 | 4.921 |
| **Farnesol** | 1749 | (*E*)-β-Caryophyllene | 0 | 2.558 | 4.382 | 6.489 | 9.194 | 0 | 2.815 | 4.594 | 6.954 | 10.079 |
| **Unknown SQT 5** | 2033 | (*E*)-β-Caryophyllene | 0 | 2.034 | 3.253 | 5.456 | 8.8 | 0 | 2.248 | 3.533 | 6.039 | 9.165 |

#### Supplementary table 5

Summary of the GLMMs analyzing the variation in induction (group-by-herbivory interaction) of volatile terpenes concentration in wild cotton leaves among genotypes and populations (random slopes; Likelihood ratio tests on nested models), as well as among geographic regions and between chemotypes (fixed effects; Wald chi-squared tests). Because they were partially nested, the effects of chemotype and region were analysed in separate models. Chemotype was included as fixed effect when testing variation among populations and genotypes. Chi-squared ($\chi^2$) values are shown, and statistically significant ($p < 0.05$) effects are indicated in bold. Df = Degrees of freedom of $\chi^2$ tests. The random slope component includes a term for the variance in slopes and one for the covariance between slopes and intercepts, unless the model could not converge, and the covariance term was excluded (italicized values).

| Leaf volatiles | **Genotype x herbivory ($\chi^2$ value, df = 2)** | **Population x herbivory ($\chi^2$ value, df = 2)** | **Region x herbivory ($\chi^2$ value, df = 1)** | **Chemotype x herbivory ($\chi^2$ value, df = 1)** |
|---|---|---|---|---|
| Total terpenes | 0.35 | 1.22 | 2.88 | 0.19 |
| (E)-β-Ocimene | 5.23 | 0.71 | 2.02 | **7.69** |
| δ-Cadinene | *0.00* | 0.88 | 0.94 | 0.29 |
| β-Caryophyllene | 0.02 | 0.48 | 4.00 | 0.02 |
| α-Pinene | *0.00* | 1.42 | 3.25 | 0.01 |
| γ-Terpinene | 0.04 | 0.59 | 1.35 | - |

#### Supplementary table 6

Volatiles quantified in the headspace samples collected from undamaged and damaged wild cotton plants (>10% of samples). Median emission rates (normalized by the total leaf area) and quantiles. RI: retention index. RRF: standard used for quantification by relative response factor. Compounds marked with an asterisk were authenticated via comparison to standards. The RI of the unknown aldoxime could not be measured as we had no alkane eluting before that analyte. Distinction between the two 2-methylbutyraldoximes isomers is tentative.

| | | | Undamaged plants ng $h^{-1}$ $cm^{-2}$ | | | | | Damaged plants ng $h^{-1}$ $cm^{-2}$ | | | | |
|---|---|---|---|---|---|---|---|---|---|---|---|---|
| **Compounds** | **RI** | **RRF** | **Q10** | **Q25** | **Median** | **Q75** | **Q90** | **Q10** | **Q25** | **Median** | **Q75** | **Q90** |
| **Unknown aldoxime** | - | (*Z*)-3-Hexenal | 0 | 0 | 0 | 0 | 0 | 0 | 0.009 | 0.054 | 0.1 | 0.217 |
| **(*E*)-2-Hexenal*** | 857 | (*E*)-2-Hexenal | 0 | 0 | 0 | 0.008 | 0.014 | 0 | 0.012 | 0.029 | 0.047 | 0.093 |
| **(*Z*)-3-Hexenol*** | 859 | (*Z*)-3-Hexenol | 0 | 0 | 0 | 0.005 | 0.034 | 0.005 | 0.013 | 0.02 | 0.056 | 0.176 |
| **(E)-2-Methylbutyraldoxime*** | 862 | (*E*)-2-Hexenol | 0 | 0 | 0 | 0 | 0 | 0 | 0.017 | 0.081 | 0.251 | 0.517 |
| **(*Z*)-2-Methylbutyraldoxime*** | 868 | (*E*)-2-Hexenol | 0 | 0 | 0 | 0 | 0 | 0 | 0.007 | 0.028 | 0.074 | 0.156 |
| **(*E*)-2-Hexenol*** | 871 | (*E*)-2-Hexenol | 0 | 0 | 0 | 0.004 | 0.008 | 0 | 0.005 | 0.015 | 0.029 | 0.046 |
| **2-Methylbutyl acetate** | 881 | (*Z*)-3-Hexenyl acetate | 0 | 0 | 0 | 0 | 0.005 | 0.014 | 0.023 | 0.053 | 0.138 | 0.315 |
| **(*Z*)-2-Pentenyl acetate** | 917 | (*Z*)-3-Hexenyl acetate | 0 | 0 | 0 | 0.001 | 0.006 | 0 | 0.002 | 0.011 | 0.023 | 0.049 |
| **α-Thujene** | 928 | α-Pinene | 0 | 0 | 0 | 0 | 0.005 | 0 | 0 | 0.003 | 0.008 | 0.021 |
| **α-Pinene*** | 934 | α-Pinene | 0.027 | 0.059 | 0.117 | 0.241 | 0.471 | 0.183 | 0.335 | 0.504 | 0.863 | 1.147 |
| **Methyl (*Z*)-3-hexenoate** | 936 | (*Z*)-3-Hexenyl acetate | 0 | 0 | 0 | 0 | 0 | 0 | 0 | 0.005 | 0.069 | 0.263 |
| **Camphene** | 949 | α-Pinene | 0 | 0 | 0.005 | 0.008 | 0.018 | 0.006 | 0.012 | 0.019 | 0.033 | 0.065 |
| **Methyl (*E*)-2-hexenoate** | 969 | (*Z*)-3-Hexenyl acetate | 0 | 0 | 0 | 0 | 0 | 0 | 0 | 0 | 0.02 | 0.05 |
| **β-Pinene*** | 977 | α-Pinene | 0 | 0.007 | 0.023 | 0.049 | 0.096 | 0.031 | 0.061 | 0.095 | 0.159 | 0.224 |
| **β-Myrcene*** | 992 | β-Myrcene | 0.002 | 0.016 | 0.062 | 0.146 | 0.279 | 0.05 | 0.084 | 0.165 | 0.329 | 0.514 |
| **(*Z*)-3-Hexenyl acetate*** | 1008 | (*Z*)-3-Hexenyl acetate | 0 | 0 | 0.009 | 0.029 | 0.063 | 0.069 | 0.11 | 0.22 | 0.489 | 1.023 |
| **n-Hexyl acetate*** | 1015 | (*Z*)-3-Hexenyl acetate | 0 | 0 | 0 | 0 | 0.003 | 0 | 0 | 0.004 | 0.011 | 0.02 |
| **(*Z*)-2-Hexenyl acetate** | 1018 | (*Z*)-3-Hexenyl acetate | 0 | 0 | 0 | 0.004 | 0.006 | 0 | 0 | 0.008 | 0.013 | 0.021 |
| **p-Cymene** | 1026 | α-Pinene | 0 | 0 | 0 | 0.009 | 0.02 | 0 | 0 | 0 | 0.017 | 0.029 |
| **Limonene*** | 1030 | α-Pinene | 0 | 0 | 0.006 | 0.024 | 0.052 | 0 | 0.012 | 0.028 | 0.059 | 0.104 |
| **(*E*)-β-Ocimene*** | 1049 | (*E*)-β-Ocimene | 0 | 0 | 0.005 | 0.023 | 0.05 | 0.088 | 0.141 | 0.249 | 0.66 | 1.279 |
| **γ-Terpinene*** | 1060 | α-Pinene | 0 | 0 | 0.003 | 0.033 | 0.121 | 0 | 0.002 | 0.009 | 0.117 | 0.277 |
| **(*Z*)-Linalool oxide** | 1074 | Linalool | 0 | 0 | 0 | 0 | 0.004 | 0.032 | 0.048 | 0.066 | 0.106 | 0.147 |
| **(*E*)-Linalool oxide** | 1090 | Linalool | 0 | 0 | 0 | 0.003 | 0.007 | 0 | 0 | 0.012 | 0.03 | 0.077 |
| **Methyl benzoate*** | 1097 | Benzaldehyde | 0 | 0 | 0 | 0 | 0.005 | 0.002 | 0.006 | 0.014 | 0.033 | 0.059 |
| **Linalool*** | 1100 | Linalool | 0 | 0 | 0 | 0.005 | 0.009 | 0.021 | 0.032 | 0.062 | 0.1 | 0.143 |

#### Supplementary table 6 (cont.)

| | | | Undamaged plants ng h$^{-1}$ cm$^{-2}$ | | | | | Damaged plants ng h$^{-1}$ cm$^{-2}$ | | | | |
|---|---|---|---|---|---|---|---|---|---|---|---|---|
| **Compounds** | RI | RFF | **Q10** | **Q25** | **Median** | **Q75** | **Q90** | **Q10** | **Q25** | **Median** | **Q75** | **Q90** |
| **2-Phenylethanol*** | 1115 | Benzaldehyde | 0 | 0 | 0 | 0 | 0.001 | 0 | 0 | 0.004 | 0.012 | 0.023 |
| **DMNT** | 1116 | Linalool | 0 | 0 | 0.014 | 0.043 | 0.093 | 0.144 | 0.2 | 0.394 | 0.649 | 1.049 |
| **Benzyl nitrile** | 1140 | Benzaldehyde | 0 | 0 | 0 | 0 | 0 | 0 | 0.002 | 0.009 | 0.022 | 0.041 |
| **(*E*)-Myroxide** | 1142 | (*E*)-β-Ocimene | 0 | 0 | 0 | 0 | 0 | 0 | 0 | 0.014 | 0.025 | 0.06 |
| **(*Z*)-3-Hexenyl isobutyrate** | 1143 | (*Z*)-3-Hexenyl acetate | 0 | 0 | 0 | 0 | 0.001 | 0 | 0 | 0.005 | 0.013 | 0.036 |
| **Unknown nitrile** | 1184 | Benzaldehyde | 0 | 0 | 0 | 0 | 0 | 0 | 0 | 0.052 | 0.122 | 0.216 |
| **(*E*)-3-Hexenyl butyrate** | 1187 | (*Z*)-3-Hexenyl acetate | 0 | 0 | 0 | 0 | 0 | 0.005 | 0.009 | 0.023 | 0.067 | 0.128 |
| **(*E*)-2-Hexenyl butyrate** | 1195 | (*Z*)-3-Hexenyl acetate | 0 | 0 | 0 | 0 | 0 | 0 | 0 | 0.006 | 0.021 | 0.044 |
| **Methyl salicylate*** | 1197 | Methyl salicylate | 0 | 0 | 0 | 0.007 | 0.014 | 0 | 0.002 | 0.011 | 0.028 | 0.076 |
| **(*Z*)-3-Hexenyl 2-methylbutyrate** | 1232 | (*Z*)-3-Hexenyl acetate | 0 | 0 | 0 | 0 | 0 | 0.005 | 0.009 | 0.03 | 0.079 | 0.194 |
| **(*E*)-2-Hexenyl 2-methylbutyrate** | 1238 | (*Z*)-3-Hexenyl acetate | 0 | 0 | 0 | 0 | 0 | 0 | 0.008 | 0.024 | 0.051 | 0.092 |
| **Indole*** | 1296 | Indole | 0 | 0 | 0 | 0 | 0 | 0.135 | 0.265 | 0.576 | 1.556 | 3.021 |
| **Unknown SQT 1** | 1331 | (*E*)-β-Caryophyllene | 0 | 0 | 0 | 0 | 0.002 | 0 | 0 | 0.003 | 0.008 | 0.015 |
| **Unknown SQT 2** | 1341 | (*E*)-β-Caryophyllene | 0 | 0 | 0.011 | 0.033 | 0.044 | 0.013 | 0.034 | 0.057 | 0.119 | 0.187 |
| **Methyl anthranilate*** | 1344 | Methyl salicylate | 0 | 0 | 0 | 0 | 0 | 0 | 0 | 0.006 | 0.014 | 0.036 |
| **α-Copaene*** | 1381 | (*E*)-β-Caryophyllene | 0 | 0 | 0 | 0.003 | 0.006 | 0.003 | 0.011 | 0.022 | 0.043 | 0.09 |
| **β-Elemene** | 1396 | (*E*)-β-Caryophyllene | 0 | 0 | 0 | 0 | 0 | 0 | 0 | 0 | 0.008 | 0.013 |
| **(*Z*)-Jasmone*** | 1402 | (*Z*)-Jasmone | 0 | 0 | 0 | 0 | 0 | 0 | 0 | 0.029 | 0.085 | 0.174 |
| **(*E*)-β-Caryophyllene*** | 1426 | (*E*)-β-Caryophyllene | 0 | 0.032 | 0.142 | 0.278 | 0.521 | 0.145 | 0.286 | 0.389 | 0.605 | 1.163 |
| **(*E*)-β-Farnesene*** | 1459 | (*E*)-β-Farnesene | 0 | 0 | 0 | 0 | 0 | 0 | 0 | 0.013 | 0.129 | 0.28 |
| **α-Humulene*** | 1460 | α-Humulene | 0 | 0.004 | 0.031 | 0.076 | 0.126 | 0.03 | 0.059 | 0.106 | 0.163 | 0.314 |
| **Alloaromadendrene** | 1467 | α-Humulene | 0 | 0 | 0 | 0.002 | 0.009 | 0 | 0.004 | 0.014 | 0.02 | 0.038 |
| **Bicyclogermacrene** | 1502 | α-Humulene | 0 | 0 | 0.019 | 0.088 | 0.138 | 0.037 | 0.078 | 0.17 | 0.338 | 0.547 |
| **(*E*,*E*)-α-Farnesene** | 1510 | (*E*)-β-Farnesene | 0 | 0 | 0 | 0 | 0 | 0 | 0.01 | 0.055 | 0.208 | 0.578 |
| **δ-Cadinene** | 1528 | α-Humulene | 0 | 0 | 0 | 0 | 0.004 | 0.002 | 0.008 | 0.013 | 0.028 | 0.038 |
| **TMTT** | 1580 | Geranyl acetate | 0 | 0 | 0.002 | 0.016 | 0.082 | 0.007 | 0.029 | 0.084 | 0.282 | 0.417 |
| **Spathulenol** | 1584 | (*E*)-β-Caryophyllene | 0 | 0 | 0 | 0.003 | 0.008 | 0 | 0.002 | 0.009 | 0.018 | 0.029 |
| **Caryophyllene oxide** | 1590 | (*E*)-β-Caryophyllene | 0 | 0 | 0 | 0.004 | 0.016 | 0 | 0 | 0.006 | 0.011 | 0.02 |

#### Supplementary table 7

Summary of the GLMMs analysing the variation in HIPV emission rates from damaged wild cotton plants among populations (random intercept; Likelihood ratio tests on nested models), as well as among geographic regions and between chemotypes (fixed effects; Wald chi-squared tests). Because they were partially nested, the effects of chemotype and region were analysed in separate models. Chemotype was included as fixed effect when testing variation among populations. Damage intensity was controlled for in all models. Chi-squared (χ 2) values are shown, and statistically significant ($p < 0.05$) effects are indicated in bold. Df = Degrees of freedom. Ce: Celestún, Si: Sisal, Ch: Chixculub, Co: Coloradas; A: chemotype A. B: chemotype B.

| HIPV emitted | **Population ($\chi^2$ value, df = 1)** | **Region ($\chi^2$ value, df = 3)** | **Effect** | **Chemotype ($\chi^2$ value, df = 1)** | **Effect** | **Figure** |
|---|---|---|---|---|---|---|
| Total HIPVs | 1.34 | 2.89 | - | 0.00 | - | Fig. S10 |
| (*E*,*E*)-α-Farnesene | **4.23** | **25.45** | Ch < Co ≤ Ce ≤ Si | 0.72 | - | Fig. 6 |
| (*E*)-Myroxide | **21.88** | **11.44** | Ch ≤ Co ≤ Ce ≤ Si | 0.04 | - | Fig. S10 |
| (*E*)-Methylbutyraldoxime | **19.25** | **62.02** | Co ≤ Ch < Ce ≤ Si | **7.24** | B > A | Fig. 6 |
| (Z)-Methylbutyraldoxime | **20.86** | **71.29** | Co ≤ Ch < Ce ≤ Si | 3.50 | - | Fig. S10 |
| Unknown oxime | 1.89 | **25.79** | Co ≤ Ch < Ce ≤ Si | **5.07** | B > A | Fig. 6 |
| Benzyl nitrile | 0.11 | **8.83** | Ch ≤ Co ≤ Si ≤ Ce | **4.71** | B > A | Fig. 6 |
| Unknown nitrile | 1.55 | **8.03** | Ce ≤ Si ≤ Ch ≤ Co | **7.10** | A > B | Fig. 6 |
